# A sensitive and specific genetically encodable biosensor for potassium ions

**DOI:** 10.1101/2021.10.07.463410

**Authors:** Sheng-Yi Wu, Yurong Wen, Nelson Bernard Calixte Serre, Cathrine Charlotte Heiede Laursen, Andrea Grostøl Dietz, Brian R. Taylor, Abhi Aggarwal, Vladimir Rancic, Michael Becker, Klaus Ballanyi, Kaspar Podgorski, Hajime Hirase, Maiken Nedergaard, Matyáš Fendrych, M. Joanne Lemieux, Daniel F. Eberl, Alan R. Kay, Robert E. Campbell, Yi Shen

## Abstract

Potassium ions (K^+^) play a critical role as an essential electrolyte in all biological systems. Here we report the crystal structure-guided optimization and directed evolution of an improved genetically encoded fluorescent K^+^ biosensor, GINKO2. GINKO2 is highly sensitive and specific for K^+^ and enables *in vivo* detection of K^+^ dynamics in multiple species.

## Main

The potassium ion (K^+^) is one of the most abundant cations across biological systems^1^. It is involved in a variety of cellular activities in organisms ranging from prokaryotes to multicellular eukaryotes^2,3^. While studies of other biologically important cations, notably calcium ion (Ca^2+^), have been revolutionized by the availability of high-performance genetically encoded biosensors^4,5^, the development of analogous biosensors for K^+^ has lagged far behind. Currently, the primary methods to monitor K^+^ are electrode-based methods, which are low throughput and not practical for subcellular level detection^6^, and synthetic indicator dyes^7^, which require invasive loading and washing procedures, and discriminate poorly between Na^+^ and K^+^.

A high-performance genetically encoded fluorescent biosensor for K^+^ could enable a variety of applications that are currently challenging, by enabling targeted expression and non-invasive *in vivo* imaging. We have previously reported a prototype intensiometric K^+^ biosensor, designated GINKO1, based on the insertion of K^+^-binding protein (Kbp) into enhanced green fluorescent protein (EGFP)^8^. Ratiometric genetically encoded biosensors have also been reported^8,9^. To create a more robust K^+^ biosensor with broader utility, we undertook the effort to further improve the sensitivity and specificity of GINKO1.

To better understand the K^+^-dependent fluorescence response mechanism of GINKO1 and facilitate further engineering, we determined the crystallographic structure of GINKO1 in the K^+^-bound state at 1.85 Å (**Fig. 1a, Supplementary Table 1, Supplementary Fig. 1, Supplementary Note**). Well-diffracting crystals of the unbound state were unattainable. The K^+^-bound crystal structure revealed the location and coordination geometry of the K^+^ binding site of Kbp (**Fig. 1b**), which was not apparent from the previously reported NMR structure (**Supplementary Fig. 2**)^10^. Notably, the backbone carbonyl oxygen atoms of six amino acids (V154, K155, A157, G222, I224, and I227) coordinate the ion, similar to the coordination modes of K^+^ selectivity filters in K^+^ channel KcsA and K^+^ transport protein TrkH^11^.

**Figure 1.**
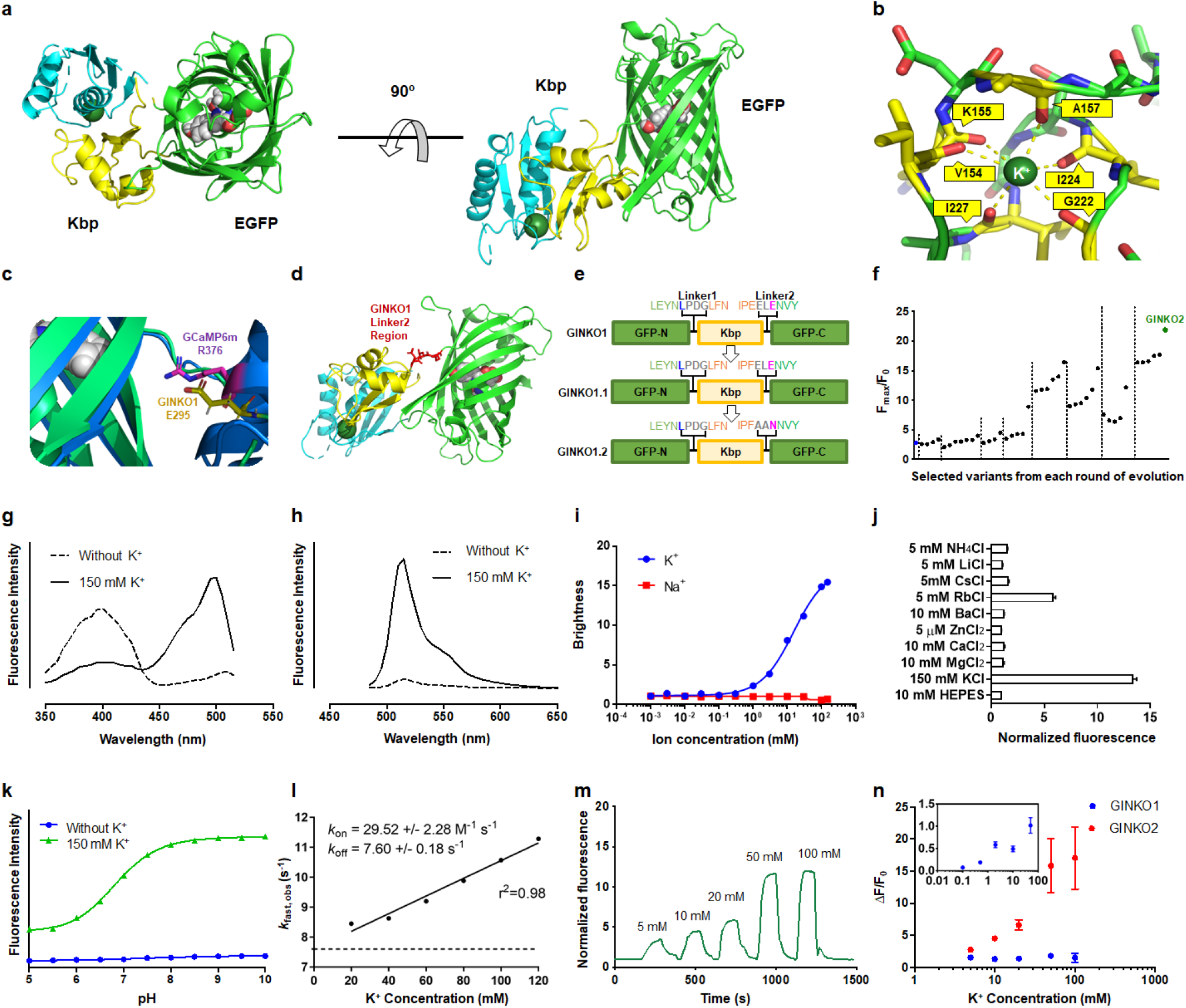
GINKO1 structure and GINKO2 engineering and characterization. **a** Cartoon representation of the structure of GINKO1 with the BON (bacterial OsmY and nodulation) domain of Kbp in cyan, the LysM (lysin motif) domain of Kbp in yellow, and the EGFP in green. The chromophore and the K^+^ ion (green) are shown in sphere representation. **b** The K^+^ is coordinated by carbonyl backbone atoms of six amino acids. **c** Crystal structure alignment of GINKO1 and GCaMP6m. Alignment of R376 (magenta sticks) of GCaMP6m (PDB: 3WLC) to E295 (yellow sticks) of GINKO1. GINKO1 is represented by green ribbons and GCaMP6m is represented by blue ribbons. Both residues point towards the chromophore of the EGFP (sphere representation). **d** GINKO1 linker regions. Linker 2 region is highlighted in red. **e** Structure-guided optimization of GINKO. Amino acid sequences of linker regions of GINKO1, GINKO1.1, and GINKO1.2 are labeled. **f** Selected variants in directed evolution of GINKO. Each dot represents a variant that was selected for its improved F_max_/F_0_ in the lysate screening. The final variant GINKO2 is highlighted in green. **g** Excitation spectra for GINKO2. **H** Emission spectra for GINKO2. **i** K^+^ and Na^+^ titration of GINKO2. **j** Ion specificity of GINKO2 (n = 3). The concentrations of cations used were above their physiological concentrations. **k** pH titrations of GINKO2. **l** Kinetics of GINKO2 (n = 3). **m** Representative *in situ* K^+^ titration with digitonin-permeabilized HeLa cells. **n** GINKO1 (n = 6) and GINKO2 (n = 10) response curves based on *in situ* titration in HeLa cells. GINKO1 response curve from 0.1 to 50 mM K^+^ is shown in the inset (n = 17).

Structure-guided mutagenesis and directed evolution were used to optimize GINKO1. Alignment of the structure of GINKO1 with that of GCaMP6 (**Fig. 1c**)^12^ revealed that GINKO1 E295 structurally aligns with GCaMP6 R376. R376 is engaged in a water-mediated interaction with the chromophore in GCaMP, likely acting to communicate the Ca^2+^-dependent conformational change in the Ca^2+^-binding domain to the GFP chromophore^13^. We thus mutated GINKO E295 to basic and hydrophobic amino acids (K/R/W/Y/P/L/F), with the hypothesis that these residues could similarly modulate the chromophore environment by introducing an opposite charge or removing the charge altogether. Among the E295 mutants, E295F was found to have the largest intensity change F_max_/F_0_ of 3.0, thus it was selected as GINKO1.1 (**Supplementary Fig. 3**). Structural and mechanistic analysis of high-performance biosensors suggested that the linker regions are of particular interest for biosensor optimization^14^, therefore we randomized the linker residues connecting EGFP to Kbp and screened for larger F_max_/F_0_ (**Fig. 1d**). This yielded GINKO1.2 with a linker sequence of A296/A297/N298 (**Fig. 1e**), which has a 30% improvement in F_max_/F_0_. We further optimized GINKO via directed evolution in *E. coli* (**Supplementary Fig. 4**). After multiple rounds of iterative evolution, we identified the best variant measured, designated GINKO2, with improved brightness and K^+^ response (**Fig. 1f, Supplementary Table 2, Supplementary Fig. 5, Supplementary Note**).

To characterize GINKO2 *in vitro*, we determined its fluorescence spectra, brightness, affinity, fluorescence change (F_max_/F_0_), specificity, kinetics, and pH dependence. Upon K^+^ binding, GINKO2 exhibits a ratiometric change in excitation spectrum (R_max_/R_0_ = 21.0 ± 0.4, where R represents the excitation ratio of 500 nm/400 nm), enabling ratiometric detection of K^+^ concentration (**Fig. 1g**). GINKO2 emission exhibited a 15x intensiometric increase at its peak of 515 nm (**Fig. 1h**). GINKO2 has a high brightness of 15.5 mM^-1^cm^-1^ in the K^+^-bound state, a 1.8x improvement over GINKO1 (8.5 mM^-1^cm^-1^) (**Supplementary Table 3**). The affinity (*K*_d_) of purified GINKO2 for K^+^ is 15.3 mM. While GINKO1 shows substantial sodium (Na^+^)-dependent fluorescence response at concentrations below 150 mM, preventing its application when Na^+^ is abundant^8^, GINKO2 is not responsive to Na^+^ at concentrations up to 150 mM, thus showing an improved specificity (**Fig. 1i**). GINKO2 responds to Rb^+^, which has a similar ionic radius to K^+^, but does not respond to Zn^2+^, Mg^2+^, Ca^2+^, Ba^2+^, Cs^+^, Li^+^, or NH_4_^+^ (**Fig. 1j, Supplementary Fig. 6, Supplementary Fig. 7**). Having low physiological concentrations (∼0.1 mM)^15^, Rb^+^ is unlikely to interfere with GINKO2 biosensing, except when used as a substitute for K^+^ in experimental settings. Kinetic measurements revealed a *k*_on_ of 29.5 ± 2.3 M^-1^s^-1^ and *k*_off_ of 7.6 ± 0.2 s^-1^ (**Fig. 1l**). GINKO2 is sensitive to pH with a p*K*_a_ of 6.8 in the K^+^-bound state (**Fig. 1k**), which necessitates consideration of the pH environment in its applications (**Supplementary Note**). In permeabilized HeLa cells, GINKO2 showed a ΔF/F_0_ of 17 when titrated with 5–100 mM K^+^, substantially larger than that of GINKO1 (ΔF/F_0_ = 1.5) (**Fig. 1m, 1n**). Overall, GINKO2 displays superior sensitivity and specificity over GINKO1.

To determine whether GINKO2 could be used to monitor intracellular K^+^ in bacteria, we attempted to use it to monitor the decreasing intracellular K^+^ concentration in *E. coli* that can be induced by growth in low-K^+^ medium. Real-time imaging of intracellular K^+^ concentration dynamics could allow us to establish the relations between extracellular low K^+^ input, intracellular K^+^ availability, and bacterial growth. The excitation ratiometric change of GINKO2 presents a unique solution to monitor K^+^ concentration changes in proliferating *E. coli*, in which intensity-based measurements are impeded by the increasing biosensor expression level. GINKO2-expressing *E. coli* growing in medium with 20 µM K^+^ exhibited a 58% decrease in excitation ratio R_470/390_ (**Fig. 2a**), corresponding to an estimated decrease in intracellular K^+^ from 100 ± 21 mM to 28 ± 4 mM (**Supplementary Fig. 8**). In contrast, cells growing in a medium with 800 µM K^+^ showed unchanged intracellular K^+^ concentration during the same growth period (**Fig. 2a**). An excitation ratiometric pH biosensor pHluorin^16^ was used to confirm that the intracellular pH remained stable. This application of GINKO2 demonstrated its practicality for real-time monitoring of intracellular K^+^ in *E. coli*.

**Figure 2.**
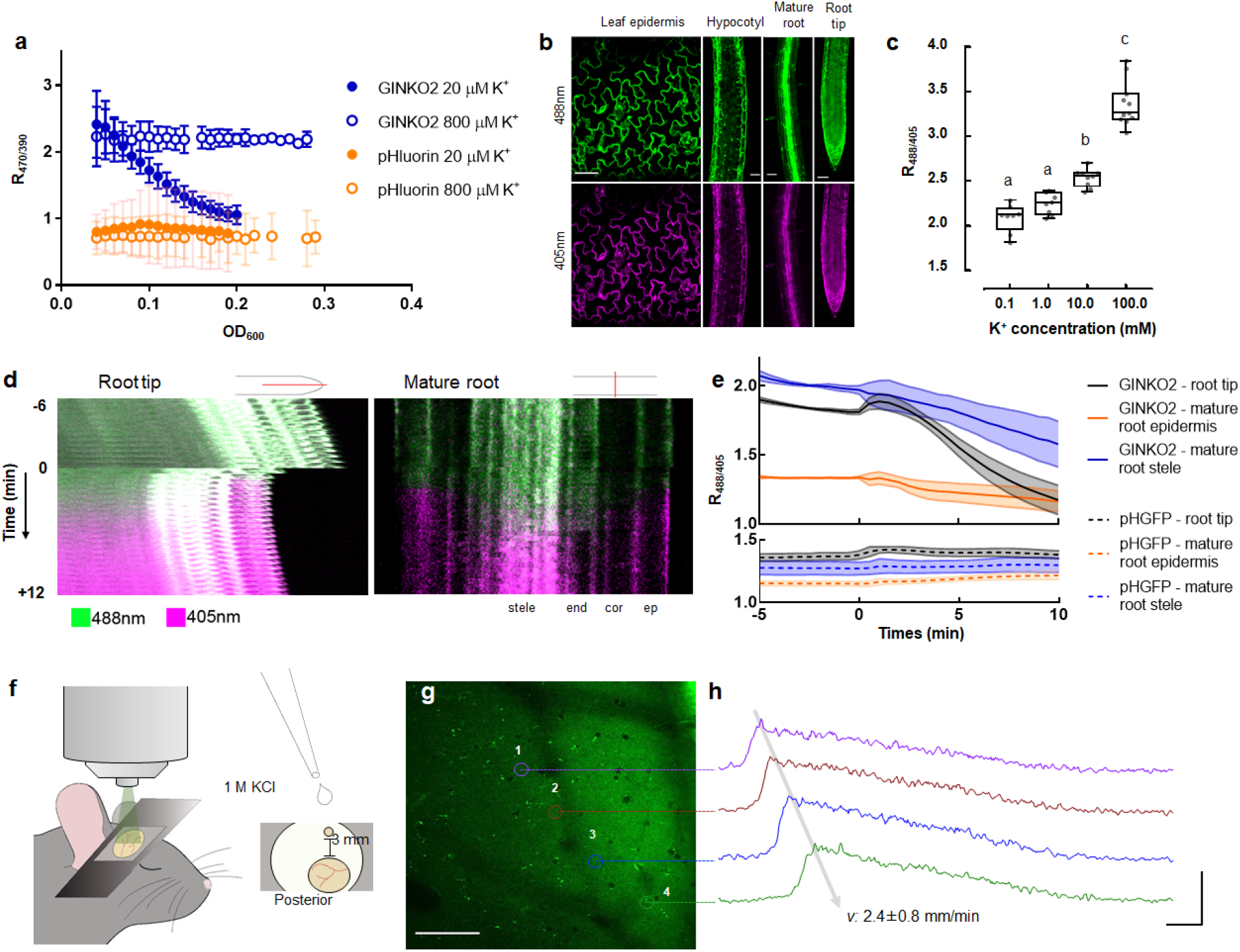
Applications of GINKO2 in multiple model organisms. GINKO2 was applied in *E. coli* (**a**), *Arabidopsis thaliana* (**b**–**e**), and mice (**f**–**h**). **a** Excitation ratio (R_470/390_) of GINKO2 in *E. coli* cells growing in K^+^-deficient media. Optical density at 600 nm (OD_600_) reflects cell density during the growth. Two low K^+^ concentrations (Open circle: 800 µM, solid circle: 20 µM) were used for the experiment: only the medium supplemented with 20 µM K^+^ induced detectable K^+^ decrease during the growth. n = 6–8 for *E. coli* expressing GINKO2 in 20 μM K^+^; n = 3–8 for *E. coli* expressing GINKO2 in 800 μM K^+^; n = 3 for *E. coli* expressing pHluorin in 20 μM K^+^; n = 3–6 for *E. coli* expressing pHluorin in 800 μM K^+^. **b** Expression and characterization of GINKO2 in *Arabidopsis thaliana*. Representative fluorescence images of GINKO2 expression under the constitutive g10-90 promoter in tissues excited at 405 nm and 488 nm. Scale bar = 50 µm. **c** Effect of increasing concentrations of KCl and 2 µM valinomycin on g10-90::GINKO2 R_488/405_ after 6 hours of K^+^ depletion with a 0 mM KCl and 2 µM valinomycin pre-treatment. n = 16-21 individual seedlings. Letters indicate the significantly different statistical groups with P < 0.05 minimum. Statistical analysis was conducted with a nonparametric multiple comparison **d** Representative timelapse of g10-90::GINKO2 fluorescences after treatment with 100 mM NaCl at time zero in the root tip and the mature root depleted with a 0 mM KCl and 2 µM valinomycin pre-treatment. The location of the selection is indicated in red above the pictures. Both channels are represented as a composite image. ep: epidermis, cor: cortex, end: endodermis. The root tip shrank upon the NaCl application due to the osmotic pressure change. **e** Effect of 100 mM NaCl on g10-90::GINKO2 R_488/405_ (top panel) in root tips, mature root stele, and epidermis with K^+^ depleted for six hours. Treatment was applied at time zero. n = 14 (root tip), 8 (mature root stele and epidermis) individual seedlings. pHGFP (bottom panel), a pH biosensor modified from ratiometric pHluorin for plant expression^23^, was used as controls. n = 9 individual seedlings for root tips, mature root stele and epidermis. The fluorescence was measured on the visible areas of the tissues shown in **d. f** Experimental setup of 2-photon microscopy in anesthetized mice. Cortical spreading depression (CSD) was induced using 1 M KCl applied to a separate frontal craniotomy (small circle) of the imaging window (large circle) at a distance of 3 mm. **g** Averaged image of GINKO2 in the somatosensory cortex (−70 µm) obtained using 2-photon microscopy. GINKO2 was applied by bath application one hour before imaging. The image depicts the regions of interest (ROIs) corresponding to traces in (**h**). Scale bar 100 µm. **h** Example of traces from ROIs in the same animal, depicting the first CSD wave. x-axis: 5 sec, y-axis: 100% ΔF/F, mean ± s.d.

To evaluate GINKO2 utility *in vivo* in plants, we sought to use GINKO2 to monitor intracellular K^+^ concentration changes in *Arabidopsis thaliana* under stress conditions. K^+^ is an essential nutrient for plants and regulates root growth, drought resistance, and salt tolerance^17,18^. Despite the importance of K^+^, its detailed physiological roles and dynamics remain elusive in plants, largely due to the lack of high-performance imaging probes. *Arabidopsis thaliana* stably transformed with g10-90::GINKO2 exhibited homogeneous fluorescence in leaf epidermis, hypocotyls, primary root tips, and primary mature roots (**Fig. 2b**). GINKO2 fluorescence is visible in the cytoplasm (pH = 7.5) but absent in vacuoles (pH = 5.5)^19^. GINKO2 expression did not affect root elongation (**Supplementary Fig. 9**), morphology, or growth rate. In permeabilized and K^+^-depleted seedlings, where manipulation of intracellular K^+^ is more feasible due to the reduced vacuolar K^+^ buffering capacity (**Supplementary Note, Supplementary Fig. 10**), GINKO2 excitation ratio R_488/405_ correlated well with the medium K^+^ concentrations in the physiological range of 1–100 mM (**Fig. 2c**). We then imaged K^+^ dynamics in roots under salt (NaCl) stress. The Na^+^ influx to the roots triggers K^+^ efflux to counterbalance the membrane depolarization^20^. With 100 mM NaCl treatment, GINKO2 reported a 35%, 20%, and 13% decrease in R_488/405_ in root tips, mature root stele, and mature root epidermis, respectively, in K^+^-depleted plants (**Fig. 2d, 2e, Supplementary Movie 1, 2**). These results demonstrated that GINKO2 is a suitable tool for reporting cytoplasmic K^+^ dynamics *in vivo* in plants.

To further explore GINKO2 applications, we tested whether GINKO2 is able to report extracellular K^+^ changes *in vivo* during cortical spreading depression (CSD) in the mouse brain. CSD is a propagating, self-regenerating wave of neuronal depolarization moving through the cortex, and is associated with severe brain dysfunctions such as migraine aura and seizures^21^. On the molecular level, CSD is accompanied by propagating waves of robust increases in extracellular K^+^ to a concentration of 30–80 mM, from a baseline of 3.5–5 mM^22^. ‘To evaluate GINKO2 during CSD, purified GINKO2 protein (6.55 mg/mL in aCSF) was exogenously applied to the extracellular space of deeply anesthetized mice above the somatosensory cortex (**Fig. 2f**). In response to application of 1 M KCl to a separate frontal craniotomy^22^ (**Fig. 2f**), we observed multiple waves of GINKO2 fluorescence intensity increase, propagating at a velocity of 2.4 ± 0.8 mm/min (**Fig. 2g, 2h, Supplementary Fig. 11a, Supplementary Movie 3**). The fluorescence intensity increased by 1.0 ± 0.2× (**Supplementary Fig. 11b**), with a fast rise at 29 ± 7% s^-1^ and a significantly slower decay at 0.03 ± 0.01% s^-1^ (**Supplementary Fig. 11c**). The duration of the waves (width at half maximum) was 22 ± 6 s (**Supplementary Fig. 11d**). Overall, GINKO2 is an effective tool for reporting extracellular K^+^ concentration changes *in vivo* in the mouse brain during CSD.

In conclusion, we have engineered an improved genetically encoded green fluorescent K^+^ biosensor GINKO2. Due to its excellent sensitivity and specificity, this new biosensor opens new avenues for *in vitro* and *in vivo* K^+^ imaging in a variety of tissues and model organisms.

## Methods

### Protein Engineering

pBAD and pcDNA plasmids containing the gene encoding GINKO1 were used as the templates for this work. Gene fragments and primers were ordered from Integrated DNA Technology (IDT). *E. coli* DH10B (Thermo Fisher Scientific) was used for cloning and protein expression. Site-directed mutagenesis was done with the QuikChange lightning kit (Agilent) according to the manufacturer’s instructions. Random mutagenesis was introduced via Error-prone PCR (EP-PCR). Briefly, EP-PCR was performed using Taq polymerase and the standard Taq buffer (New England Biolabs) with imbalanced dNTP (0.2 mM dATP, 0.2 mM dTTP, 1 mM dGTP, and 1 mM dCTP) and modifications of MnCl_2_ and MgCl_2_ concentrations. 5.5 mM MgCl_2_ was added to the supplier’s standard reaction buffer. MnCl_2_ was added to a final concentration of 0.15 mM and 0.30 mM to generate libraries with low-frequency and high-frequency mutations. 2% (v/v) DMSO was added to stabilize the unmatched nucleotide pairs during the amplification. PCR products were purified on 1% agarose gel, digested with *Xho*I and *Hind*III (Thermo Fisher Scientific), and ligated with a similarly digested pBAD backbone vector using T4 DNA ligase (Thermo Fisher Scientific). The transformation of electrocompetent DH10B (Thermo Fisher Scientific) was performed with the ligation products and QuikChange products. About 5,000–10,000 colonies were generated for each library. 40–80 colonies with bright to medium fluorescence were picked and inoculated at 37°C overnight. Cells were pelleted down by centrifugation at > 10,000 rpm for 30 seconds, resuspended in 200 µL of 10 mM HEPES buffer, and lysed by four freeze and thaw cycles by incubating in liquid nitrogen and 42°C water bath. The lysate was centrifuged for 5 minutes. 100 µL supernatant of each sample was then transferred to a 96-well plate. The fluorescence response was read by a Safire2 microplate reader (Tecan) with excitation at 465 nm. 10 µL of 1 M KCl was then added into each well and the fluorescence measurements were repeated. The winners were selected based on the calculated fluorescence change (F_max_/F_0_) and validated in triplicates. K^+^ titrations were performed on purified variants to further verify the fluorescence change (F_max_/F_0_) and determine the *K*_d_. The winners were selected for the next round of optimization.

### Protein expression and purification

Single colony of *E. coli* DH10B expressing GINKO variants were picked from the agar plate and inoculated in a flask containing 200–500 mL of LB supplemented with 100 µg/mL ampicillin and 0.02% (w/v) L-(+)-arabinose. The cells were cultured at 200 rpm, 37°C for 16–20 hours. GINKO variants were purified as previously described^8^. Briefly, the cells were pelleted down by centrifugation at 6000 rpm for 10 minutes and lysed by sonication. The protein was purified through affinity chromatography with Ni-NTA beads. The protein-bound beads were washed with the wash buffer supplemented with 20 mM imidazole. GINKO was eluted from the beads with the elution buffer supplemented with 500 mM imidazole. The eluted protein was then buffer exchanged to 10 mM HEPES at pH 7.4 by PD-10 columns (GE Healthcare Life Sciences) following the manufacturer’s instructions.

### Crystallization and structure determination

The His-tag affinity-purified GINKO1 protein was further applied on the size exclusion chromatography Superdex200 (GE Healthcare) column pre-equilibrated with 25 mM Tris pH 7.5, 150 mM KCl buffer. The main fractions of monodisperse protein were concentrated to around 25 mg/mL for crystallization trials. Crystallization experiments were set up in sitting drop geometry with 0.5 µl protein sample equilibrating with 0.5 µl reservoir from screen kits (Hampton and Molecular Dimensions) at room temperature. The final diffraction quality crystals were grown in 0.1 M MES pH 6.0, 20% PEG6000 after several rounds of crystallization optimization (**Supplementary Fig. 1b**). For data collection, the crystals were transferred to the crystal stabilization buffer supplemented with 10–15% PEG400 or glycerol and flash-frozen in liquid nitrogen. X-ray diffraction data were collected at GM/CA@APS beamline 23IDB, using raster to identify a well-diffracting region of an inhomogeneous rod-shaped crystal, and were initially processed with the beamline supplemented software package. The X-ray diffraction data were further integrated and scaled with the XDS suite^24^. The data collection details and statistics were summarized in crystallographic **Supplementary Table 1**. The GINKO1 structure was determined with a maximum-likelihood molecular replacement program implemented in the Phaser program^25^, using structures of the GFP (6GEL) and the K^+^ binding protein (5FIM) as search models^10,26^. The linker and K^+^ density were observed after initial refinement. The missing residues manual model rebuilding and refinement were carried out with the COOT program and PHENIX suite^27,28^. The GINKO1 structure was solved at 1.85 Å in the P1 space group with the unit cell dimension a = 46.8 Å, b = 49.3 Å, c = 83.7 Å, and α = 89.96°, β = 89.97°, γ = 80.95°. The final structure model was refined to a *R*_*work*_*/R*_*free*_ value of 0.1947/0.2252. The model contained two GINKO1 molecules each occupying one K^+^, and 892 water molecules in the asymmetric unit cell.

### *In vitro* characterization

The purified GINKO variants were titrated with K^+^ and Na^+^ to determine the fluorescence change F_max_/F_0_ and the affinity. The titration buffers were prepared in 10 mM HEPES at pH 7.4 supplemented with 0.001, 0.003, 0.01, 0.03, 0.1, 0.3, 1, 3, 10, 30, 100, and 150 mM KCl or NaCl. The buffers for specificity tests were prepared in 10 mM HEPES at pH 7.4. The buffers used for pH titrations were 10 mM HEPES adjusted with NaOH or HCl to pH 5.5, 6.0, 6.5, 7.0, 7.5, 8.0, 8.5, 9.0, 9.5, and 10.0 in the presence or absence of 150 mM KCl. The fluorescence measurements were performed in a Safire2 microplate reader (Tecan). The excitation wavelength was set at 460 nm for the emission scan from 485 to 650 nm, and the emission wavelength was set at 540 nm for the excitation scan from 350 to 515 nm. The extinction coefficient (EC) and quantum yield (QY) were determined to quantify the brightness of GINKO variants as described previously^8^. Briefly, GINKO variants fluorescence was measured in 10 mM HEPES at pH 7.4 either supplemented with 150 mM KCl or free of both K^+^ and Na^+^. To determine EC, DU800 spectrophotometer (Beckman Coulter) was used to measure the absorbance and quantify the denatured chromophores at 446 nm after base denaturation with 0.5 M NaOH^29^. The QY was determined using GINKO1 as the standard. Fluorescence was measured with the Safire2 microplate reader (Tecan). Rapid kinetic measurements of the interaction between GINKO2 and K^+^ were made using SX20 stopped-flow reaction analyzer (Applied Photophysics) using fluorescence detection. The dead time of the instrument was 1.1 ms. The excitation wavelength was set at 488 nm with 2 nm bandwidth and emission was collected at 520 nm through a 10-mm path. A total of 1000 data points were collected over three replicates at increments of 0.01 s for 10 seconds. Reactions were initiated by mixing equal volumes of diluted purified GINKO2 protein in 100 mM Tris-HCl, pH 7.20 with various concentrations of KCl (20, 40, 60, 80, 100, and 120 mM) at 20°C. 100 mM Tris-HCl buffer was used as a blank.

### Mammalian cell culture and imaging

HeLa cells were cultured in Dulbecco’s Modified Eagle Medium (DMEM, Gibco) supplemented with 10% fetal bovine serum (FBS, Gibco) and 200 U/mL penicillin-streptomycin (Thermo Fisher Scientific). The HeLa cells were transfected with pcDNA-GINKO variants by TurboFect transfection reagent (Thermo Fisher Scientific) as per the manufacturer’s instructions. The transfected cells were first treated with 10 nM digitonin for about 15 minutes in the imaging buffer (1.5 mM CaCl_2_, 1.5 mM MgSO_4_, 1.25 mM NaH_2_PO_4_, 26 mM NaHCO_3_ and 10 mM D-Glucose, pH = 7.4) saturated with 95% O_2_ / 5% CO_2_. The cells were then imaged on an upright FV1000 confocal microscope (Olympus) equipped with FluoView software (Olympus) and a 20× XLUMPlanF1 water immersion objective (NA 1.0, Olympus) with a flow rate of 10 mL/min using a peristaltic pump (Watson-Marlow). GINKO variants were excited with a 488-nm laser and emission was collected in the channel from 500 to 520 nm. The perfusion buffers were prepared in imaging buffers with various K^+^ concentrations (0.1, 0.5, 2, 5, 10, 20, 50, and 100 mM). N-methyl-D-glucamine (NMDG) was supplemented to keep osmotic pressure consistent. Fluorescence images were processed in Fiji. Regions of interest (ROI) were selected manually based on areas with green fluorescence.

### *E. coli* growth in K^+^ depleted environment

*Escherichia coli* NCM3722 cells were grown in a minimal medium with 20 mM NaH_2_PO_4_, 60 mM Na_2_HPO_4_, 10 mM NaCl, 10 mM NH_4_Cl, 0.5 mM Na_2_SO_4_, 0.4% arabinose and micronutrients^30^. Micronutrients include 20 μM FeSO_4_, 500 μM MgCl_2_, 1 μM MnCl_2_·4H_2_O, 1 μM CoCl_2_·6H_2_O, 1 μM ZnSO_4_·7H_2_O, 1 μM H_24_Mo_7_N_6_O_24_·4H_2_O, 1 μM NiSO_4_·6H_2_O, 1 μM CuSO_4_·5H_2_O, 1 μM SeO_2_, 1 μM H_3_BO_4_, 1 μM CaCl_2_, and 1 μM MgCl_2_. KCl was added at 800 µM or 20 µM. Ampicillin was added at 100 µg/mL to LB medium cultures and 20 µg/mL to minimal medium cultures. Cells were picked from single colonies from LB agar plates and cultured in LB medium for 3–5 hours at 37°C in a water bath shaker at 240 rpm. Cells were then diluted 1000 times into arabinose-containing minimal medium (800 µM KCl) and grown at 37°C in a water bath shaker at 240 rpm overnight. Cells were washed once in minimal media with the same K^+^ concentration as the final condition and diluted 500× into 96-well plates with 200 µL of arabinose minimal media in each well (20 or 800 µM KCl). The 96-well plates were incubated at 37°C in a Spark Plate reader (Tecan). Every 7 minutes a loop would run with the following actions: first, the plate was shaken for 200 seconds in the “orbital” mode with an amplitude of 4.5 mm at 132 rpm; then optical density was measured at 600 nm; fluorescence was measured at two wavelength settings: excitation of 390 nm, emission 520 nm and excitation 470 nm, emission 520 nm. Background fluorescence was subtracted from wild-type NCM3722 control matched by binning into the nearest 0.01 of OD and averaging fluorescence.

### *In vivo* K^+^ imaging in plants

*Arabidopsis thaliana* ecotype Columbia 0 (Col0) was used as the wild type and background for the expression GINKO2. GINKO2 was cloned into the pUPD2 plasmid using the GoldenBraid cloning system^31^. GINKO2 was placed under the control of the strong constitutive g10-90 promoter^32^, terminated by the Rubisco terminator, and together with the BASTA selection cassette, combined into the binary pDGB1_omega1 vector. Stable transformation of *Arabidopsis thaliana* plants was achieved by the floral dip method^33^. Transformed plants were then selected by their BASTA resistance and optimal fluorescence; single-locus insertion lines were selected for further propagation until homozygous lines were established. Seeds were surface sterilized by chlorine gas for 2 hours and sown on 1% (w/v) agar (Duchefa) with ½ Murashige and Skoog (MS, containing 10 mM K^+^ and 51 μM Na^+^, Duchefa), 1 % (w/v) sucrose, adjusted to pH 5.8 with NaOH, and stratified for 2 days at 4°C. Seedlings were grown vertically for 5 days in a growth chamber with 23°C by day (16 hours), 18°C by night (8 hours), 60% humidity, and the light intensity of 120 µmol photons m^-2^s^-1^. For the KCl gradient experiments, treatments were applied by transferring the plants to 0.7% (m/v) agarose (VWR Life Sciences) with 1.5 mM MES buffers (Duchefa) supplemented with various concentrations of KCl and adjusted to pH 5.8 with NaOH. For the depletion of cellular K^+^, seedlings were transferred to a 0 mM KCl medium containing 2 µM valinomycin (P-Lab, 10 mM in DMSO) in 1.5 mM MES buffer pH 5.8 in agarose 0.7%, for 6 hours before imaging. For the KCl gradient with depletion, seedlings were treated for 30 minutes before imaging. For NaCl treatment, seedlings were transferred from solid media to custom microfluidics chips^34^. Seedlings were first imaged in a control solution (0 mM NaCl in 1.5 mM MES buffer pH 5.8) before switching to a treatment solution (100 mM NaCl in 1.5 mM MES buffer pH 5.8). A constant flow of 3 µL/minute ± 0.01 µL/min was maintained using a piezoelectric pressure controller (OBI1, Elveflow, France) coupled with micro-flow sensors (MFS2, Elveflow, France) and the dedicated Elveflow ESI software to control both recording and the flow/pressure feedback. The root elongation toxicity assay was performed by scanning Col0 and g10-90::GINKO2 seedlings growing in square plates containing 1/2MS media for 16 hours every 30 minutes with an Epson v370 perfection scanner. Root elongation was quantified with a semi-automated workflow^34^. Microscopy imaging was performed using a vertical stage Zeiss Axio Observer 7 with Zeiss Plan-Apochromat 20x/0.8, coupled to a Yokogawa CSU-W1-T2 spinning disk unit with 50 µm pinholes and equipped with a VS401 HOM1000 excitation light homogenizer (Visitron Systems). Images were acquired using the VisiView software (Visitron Systems). GINKO2 was sequentially excited with a 488 nm and 405 nm laser and the emission was filtered by a 500–550 nm bandpass filter. Signal was detected using a PRIME-95B Back-Illuminated sCMOS Camera (1200 × 1200 pixels; Photometrics). For microfluidic experiments, the fluorescence was measured using the segmented line tool with a 40 pixels width. All microscopy image analyses were conducted using the software ImageJ Fiji v1.53c^35^. Statistical analyses were performed using R software. Boxplots represent the median and the first and third quartiles, and the whiskers extend to data points <1.5 interquartile range away from the first or third quartile; all data points are shown as individual dots. We used two-sided nonparametric Tukey contrast multiple contrast tests (mctp function) with logit approximation.

### *In vivo* imaging of CSD in mice

All experiments performed at the University of Copenhagen were approved by the Danish National Animal Experiment Committee and were in accordance with European Union Regulations. Male C57BL/6J wild-type mice 8–10 weeks old (Janvier) were used for *in vivo* studies. Mice were kept under a diurnal lighting condition (12 hours light/12 hours dark) in groups of five with free access to food and water. Mice were deeply anesthetized (ketamine: 100 mg/kg, xylazine: 20 mg/kg) and fixed to a stereotaxic stage with earbars. Body temperature was maintained at 37°C with a heating pad, and eye drops were applied. A metal head plate was attached to the skull using dental acrylic cement (Fuji LUTE BC, GC Corporation, Super Bond C&B, Sun Medical). A small craniotomy for KCl application was made on the skull above the frontal cortex (AP: 1.0 mm ML: 1.2 mm). Likewise, a 3 mm diameter craniotomy for imaging was drilled above the ipsilateral somatosensory cortex (AP: -1.5 mm, ML: 2.0 mm). To prepare the window for imaging, the dura was carefully removed before sealing half the craniotomy with a thin glass coverslip (3 mm x 5 mm, thickness: 0.13 mm, Matsunami Glass) using dental cement. Two-photon imaging was performed with a B-Scope equipped with a resonant scanner (Thorlabs), a Chameleon Vision 2 laser (Coherent, wavelength 940 nm), and an Olympus objective lens (XLPlan N × 25). The filter set for the detection of the green channel was as follows: primary dichroic mirror ZT405/488/561/680-1100rpc(Chroma); secondary dichroic mirror FF562-Di03 (Semrock); emission filter: FF03-525/50(Semrock). The power after the objective lens ranged between 15 mW and 30 mW. Images were acquired at a depth of 70 µm with a frame rate of 30 Hz. Immediately after surgery, deeply anesthetized mice were moved to the imaging stage, and 150 µL of GINKO2 (6.55 mg/mL in HEPES-aCSF) was applied to the craniotomy above somatosensory cortex 60–80 min before imaging. Anesthesia level was carefully monitored and maintained during the entire course of the experiment. Cortical spreading depression was induced by applying a small drop (50 to 150 µL) of 1 M KCl solution to the frontal craniotomy. Fluorescence images were processed in Fiji. Regions of interest (ROI) were selected manually based on areas with green fluorescence. Areas with small intense elements of green fluorescence were avoided. The mean fluorescence intensity of each ROI was calculated and smoothed by a 3-point average filter in MATLAB. The example trace in **Supplementary Fig. 11a** was calculated and smoothed by a 5-point average filter in MATLAB. Thereafter, relative fluorescence changes (ΔF/F) were calculated: F was the mean intensity of the pre-CSD period and ΔF was the difference between the signal and F. Velocity was calculated for the passage of signal intensity peak. Graphpad Prism was used to create figures. The data were represented as mean ± s.d. The slope coefficient was calculated using simple linear regression in Prism 9. Shapiro-Wilk normality test and paired t-test were performed using Prism 9.

### Data Analysis

Microsoft Excel was used for data analyses of GINKO characterizations and titrations. Graphpad Prism was used to create figures. The data are represented as mean ± s.d., except for the permeabilized HeLa titration, which is represented as mean ± s.e.m.

## Acknowledgments

This work was supported by grants from the Canadian Institutes of Health Research (CIHR, FS-154310 to REC) and the Natural Sciences and Engineering Research Council of Canada (NSERC, RGPIN 2018-04364 to REC, RGPIN-2020-05514 to KB, and RGPIN-2016-06478 to MJL). SYW was supported by NSERC Canada Graduate Scholarships – Doctoral program, Alberta Innovates Technology Future (AITF) Graduate Scholarship, and the University of Alberta. YW was supported by the Alberta Parkinson Society Fellowship and National Natural Science Foundation of China (No. 31870132 and No. 82072237). This research used resources of the Advanced Photon Source (APS), a U.S. Department of Energy (DOE) Office of Science User Facility operated for the DOE Office of Science by Argonne National Laboratory under Contract No. DE-AC02-06CH11357. X-ray crystallography was performed using Beamline 23IDB at APS. GM/CA@APS has been funded by the National Cancer Institute (ACB-12002) and the National Institute of General Medical Sciences (AGM-12006, P30GM138396). Data was also collected at beamline CMCF-ID at the Canadian Light Source, a national research facility of the University of Saskatchewan, which is supported by the Canada Foundation for Innovation (CFI), the NSERC, the National Research Council (NRC), the CIHR, the Government of Saskatchewan, and the University of Saskatchewan. We thank the University of Alberta Molecular Biology Services Unit for technical assistance. NBCS and MF thank Eva Medvecká for technical support. NBCS and MF work was supported by the European Research Council (Grant No. 803048) and Charles University Primus (Grant No. PRIMUS/19/SCI/09). BRT and TH are supported by NIH grant R01GM095903. AA and KP were supported by Howard Hughes Medical Institute. ARK and DFE thank Walter Boron for helpful suggestions for the *Drosophila* experiments. ARK and DFE were supported in part by NSF grant 2037828. CCHL, AGD, HH, and MN were supported by the Novo Nordisk Foundation (NNF19OC0058058 to HH and NNF13OC0004258 to MN) and the Lundbeck Foundation (R155-2016-552 to MN and R263-2017-4062 to AGD).

## Author Contributions

SYW developed GINKO2 variants, assembled all constructs, performed directed evolution, performed protein characterization, performed live-cell imaging experiments, analyzed data, prepared figures, and wrote the manuscript. YW performed the protein crystallization, solved the structures, and prepared the figures. NBCS performed the plant experiments and prepared the figures. CCHL and AGD performed the mice experiments and prepared the figures. BRT performed the *E. coli* growth experiment. AA and KP performed the stopped-flow experiments and prepared the figures. VR performed the cell imaging experiment. MB performed X-ray crystallography. YS supervised research, analyzed data, prepared figures, and wrote the manuscript. DFE and ARK performed *Drosophila* experiments, and supervised the research. KB, HH, MN, MF, MJL, and REC supervised research. All authors contributed to the editing and proofreading of the manuscript.

## Data availability

Plasmids and DNA sequences are available via Addgene (Addgene ID 177116, 177117). The GINKO1 structure is deposited in the Protein Data Bank (PDB ID:7VCM). Other expression plasmids, data supporting the findings in this research are available from the corresponding author upon request.

## Supplementary Information

### Supplementary Note

### Supplementary Methods

### Supplementary Movies

Supplementary Movie 1. Salt stress induced K^+^ decrease in the root tip of K^+^ depleted *Arabidopsis thaliana*.

Supplementary Movie 2. Salt stress induced K^+^ decrease in the mature root of K^+^ depleted *Arabidopsis thaliana*.

Supplementary Movie 3. Real-time monitoring of CSD waves with GINKO2 in mice.

---

**Supplementary Table 1.**
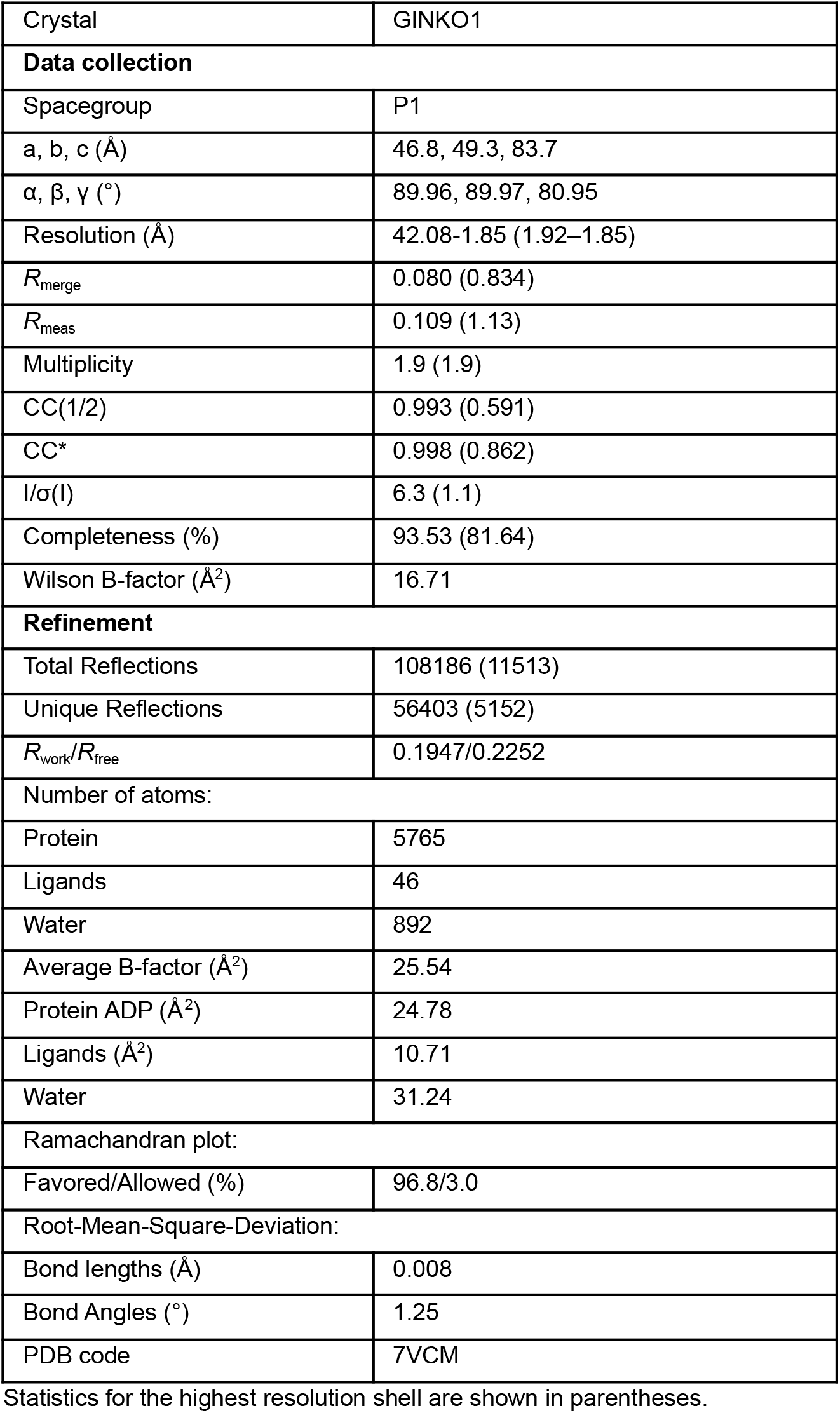
X-ray data collection and refinement statistics.

|  |  |
| --- | --- |
| Crystal | GINKO1 |
| **Data collection** |  |
| Spacegroup | P1 |
| a, b, c (Å) | 46.8, 49.3, 83.7 |
| $\alpha$, $\beta$, $\gamma$ (°) | 89.96, 89.97, 80.95 |
| Resolution (Å) | 42.08-1.85 (1.92–1.85) |
| $R_{\text{merge}}$ | 0.080 (0.834) |
| $R_{\text{meas}}$ | 0.109 (1.13) |
| Multiplicity | 1.9 (1.9) |
| CC(1/2) | 0.993 (0.591) |
| CC* | 0.998 (0.862) |
| $I/\sigma(I)$ | 6.3 (1.1) |
| Completeness (%) | 93.53 (81.64) |
| Wilson B-factor (Å<sup>2</sup>) | 16.71 |
| **Refinement** |  |
| Total Reflections | 108186 (11513) |
| Unique Reflections | 56403 (5152) |
| $R_{\text{work}}/R_{\text{free}}$ | 0.1947/0.2252 |
| Number of atoms: |  |
| Protein | 5765 |
| Ligands | 46 |
| Water | 892 |
| Average B-factor (Å<sup>2</sup>) | 25.54 |
| Protein ADP (Å<sup>2</sup>) | 24.78 |
| Ligands (Å<sup>2</sup>) | 10.71 |
| Water | 31.24 |
| Ramachandran plot: |  |
| Favored/Allowed (%) | 96.8/3.0 |
| Root-Mean-Square-Deviation: |  |
| Bond lengths (Å) | 0.008 |
| Bond Angles (°) | 1.25 |
| PDB code | 7VCM |

Statistics for the highest resolution shell are shown in parentheses.

**Supplementary Table 2.** Mutations accumulated during directed evolution.

| Variant | Mutations |
| --- | --- |
| 1 | GINKO1 Template |
| **1.2** | E295F, E296A, L297A, E298N |
| **1.3.10** | **Y40N**, **F151S**, E295F, E296A, L297A, E298N |
| 1.3.11 | **Q211L**, E295F, E296A, L297A, E298N |
| 1.3.25 | **K235E**, E295F, E296A, L297A, E298N |
| **1.5.6** | Y40N, F151S, **N152D**, E295F, E296A, L297A, E298N |
| **1.5.20** | Y40N, F151S, **K171E**, **V204L**, E295F, E296A, L297A, E298N |
| **1.5.21** | Y40N, F151S, **G167C**, **K201R**, **V261A**, **A265D**, E295F, E296A, L297A, E298N, **N299Y** |
| **1.5.36** | Y40N, F151S, **T205A**, **K259N**, E295F, E296A, L297A, E298N |
| **1.5.38** | **K102E**, **K201R**, **K235E**, E295F, E296A, L297A, E298N, **F373Y** |
| **1.5.41** | Y40N, F151S, **K284R**, E295F, E296A, L297A, E298N |
| 1.6.7 | Y40N, F151S, **V154A**, T205A, K259N, E295F, E296A, L297A, E298N |
| 1.6.8 | K102E, **D130N**, **I187F**, K201R, K235E, E295F, E296A, L297A, E298N, F373Y |
| **1.6.15** | **S31Y**, K102E, T205A, K259N, E295F, E296A, L297A, E298N |
| 1.6.40 | **V2A**, Y40N, **D130G**, F151S, N152D, E295F, E296A, L297A, E298N, **E385V** |
| **1.7.7** | S31Y, K102E, T205A, K259N, E295F, E296A, L297A, E298N, **K356R** |
| 1.7.8 | S31Y, K102E, **N194D**, T205A, K259N, E295F, E296A, L297A, E298N |
| **1.9.8** | S31Y, **I94T**, K102E, T205A, **N266D**, K259N, E295F, E296A, L297A, E298N, K356R |
| 1.9.13 | S31Y, K102E, T205A, K259N, E295F, E296A, L297A, E298N, K356R, **M383R** |
| 1.9.15 | S31Y, K102E, **D173V**, **Q174L**, T205A, K259N, E295F, E296A, L297A, E298N, **K303Q**, K356R |
| 1.9.20 | S31Y, K102E, **N183H**, T205A, K259N, E295F, E296A, L297A, E298N, **K312R**, K356R |
| 2.0.17 | S31Y, I94T, K102E, T205A, **T241A**, N266D, K259N, E295F, E296A, L297A, E298N, K356R |
| **2.0.29** | S31Y, I94T, K102E, **N183H**, T205A, N266D, K259N, E295F, E296A, L297A, E298N, K356R |
| **2.1.5** | S31Y, I94T, K102E, **K127E**, N183H, T205A, N266D, K259N, **E295Y**, E296A, L297A, E298N, K356R |
| 2.2.25<br>(**GINKO2**) | S31Y, I94T, K102E, K127E, N183H, **Q179R**, **I197V**, T205A, N266D, K259N, E295Y, E296A, L297A, E298N, **Q307E**, K356R, **K388M** |

The mutations in the template are in black and the new mutations are in red. The variants in bold were used as templates for the next round of directed evolutions. The selected variants in Library 1.4 failed to yield better variants than GINKO<sub>1.3.10</sub> and were mixed as templates for Library 1.5. Similarly, a mix of GINKO1.5 variants was used as the template to generate Library 1.6.

**Supplementary Table 3.**
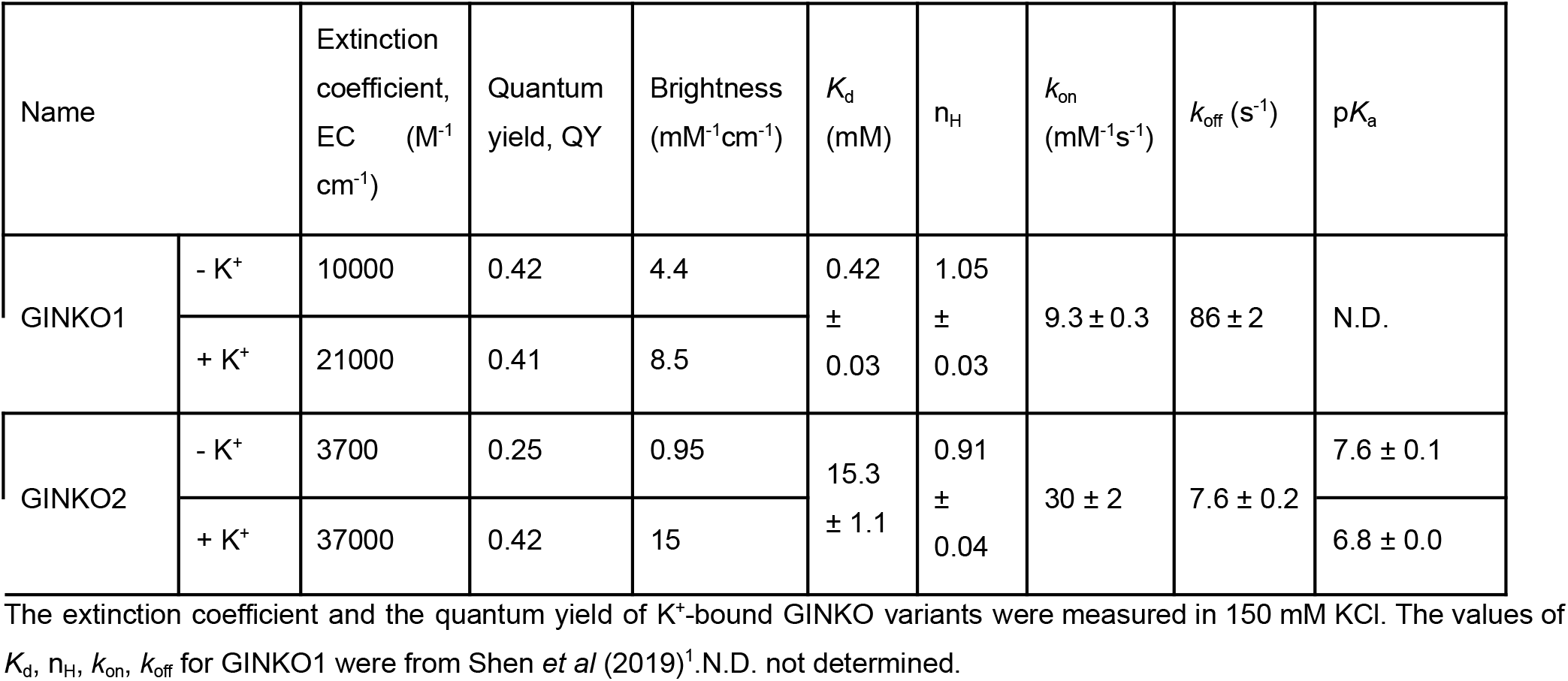
Summary of GINKO1 and GINKO2 photophysical characteristics.

| Name | | Extinction coefficient, EC (M<sup>-1</sup> cm<sup>-1</sup>) | Quantum yield, QY | Brightness (mM<sup>-1</sup>cm<sup>-1</sup>) | $K_d$ (mM) | $n_H$ | $k_{on}$ (mM<sup>-1</sup>s<sup>-1</sup>) | $k_{off}$ (s<sup>-1</sup>) | p*K<sub>a</sub>* |
| --- | --- | --- | --- | --- | --- | --- | --- | --- | --- |
| GINKO1 | - K<sup>+</sup> | 10000 | 0.42 | 4.4 | 0.42 ± 0.03 | 1.05 ± 0.03 | 9.3 ± 0.3 | 86 ± 2 | N.D. |
|  | + K<sup>+</sup> | 21000 | 0.41 | 8.5 |  |  |  |  |  |
| GINKO2 | - K<sup>+</sup> | 3700 | 0.25 | 0.95 | 15.3 ± 1.1 | 0.91 ± 0.04 | 30 ± 2 | 7.6 ± 0.2 | 7.6 ± 0.1 |
|  | + K<sup>+</sup> | 37000 | 0.42 | 15 |  |  |  |  | 6.8 ± 0.0 |

The extinction coefficient and the quantum yield of K<sup>+</sup>-bound GINKO variants were measured in 150 mM KCl. The values of $K_d$ , $n_H$ , $k_{on}$ , $k_{off}$ for GINKO1 were from Shen *et al* (2019)<sup>1</sup>. N.D. not determined.

**Supplementary Figure 1.**
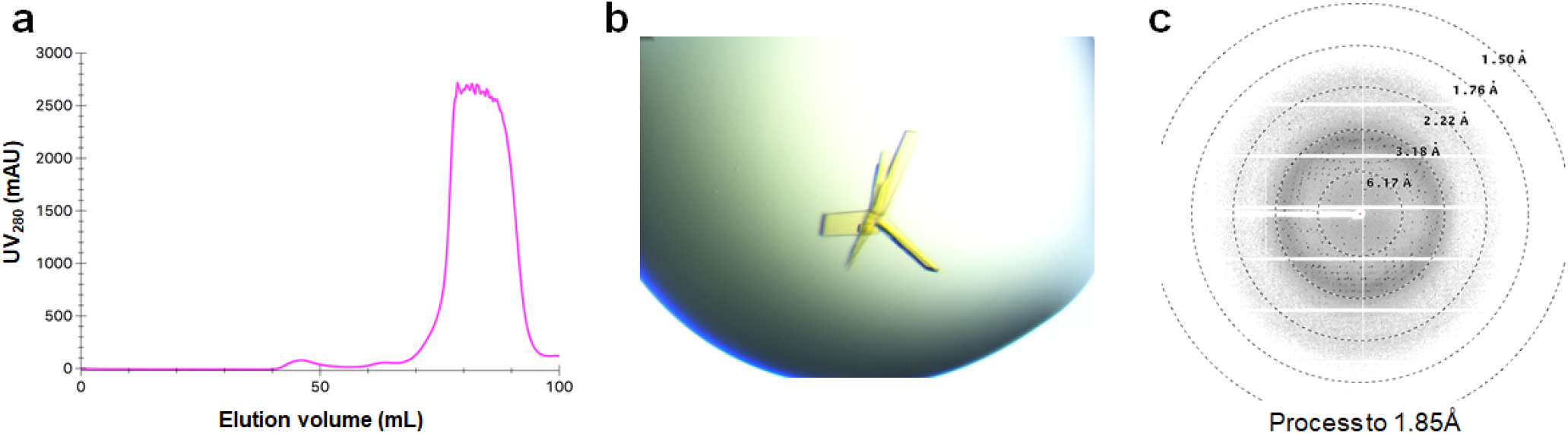
Purification, crystallization, and X-ray crystallography of GINKO1. **a** Size exclusion chromatography was used to confirm monomericity of GINKO1. **b** Image of GINKO1 crystals in TBS supplemented with 150 mM KCl. **c** The X-ray diffraction pattern of GINKO1.

**Supplementary Figure 2.**
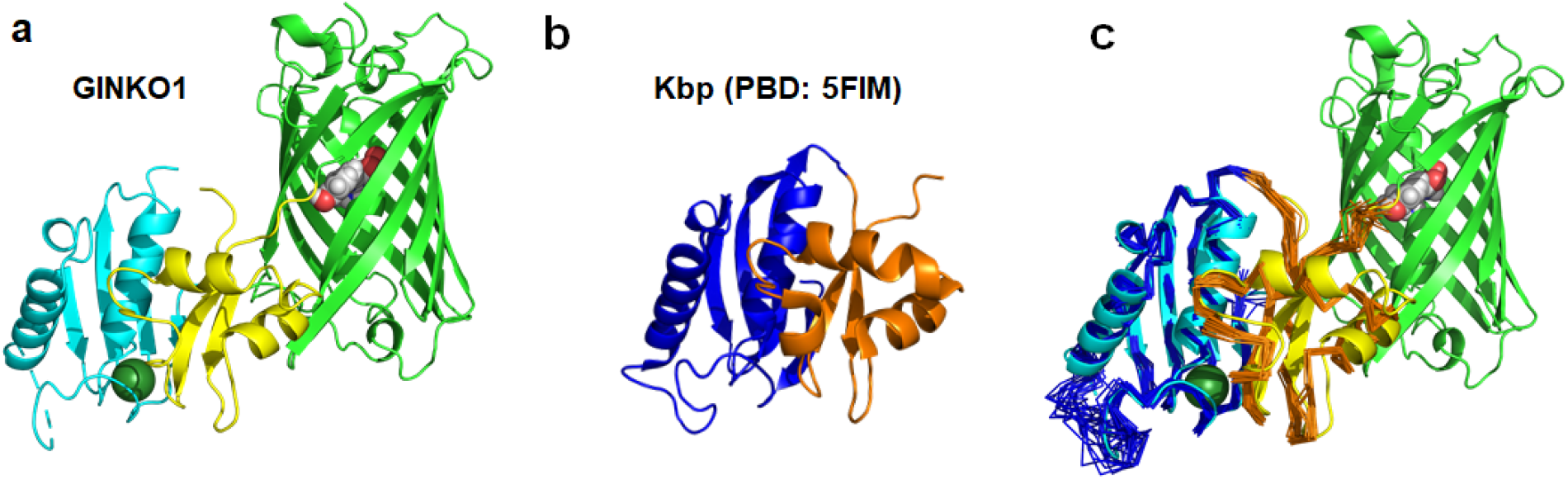
Structural comparison of GINKO1 and Kbp **a** Cartoon representation of GINKO1: EGFP domain is in green, Kbp BON domain is in cyan, and Kbp LysM domain is in yellow. The chromophore in the EGFP barrel is shown in sphere representation. The K^+^ ion is shown as a green sphere. **b** Cartoon representation of the NMR structure of Kbp (PDB ID: 5FIM). BON domain is in blue, LysM domain is in orange. **c** Structure alignment of GINKO1 and Kbp. Kbp is shown in ribbon representation with all 20 statuses.

**Supplementary Figure 3.**
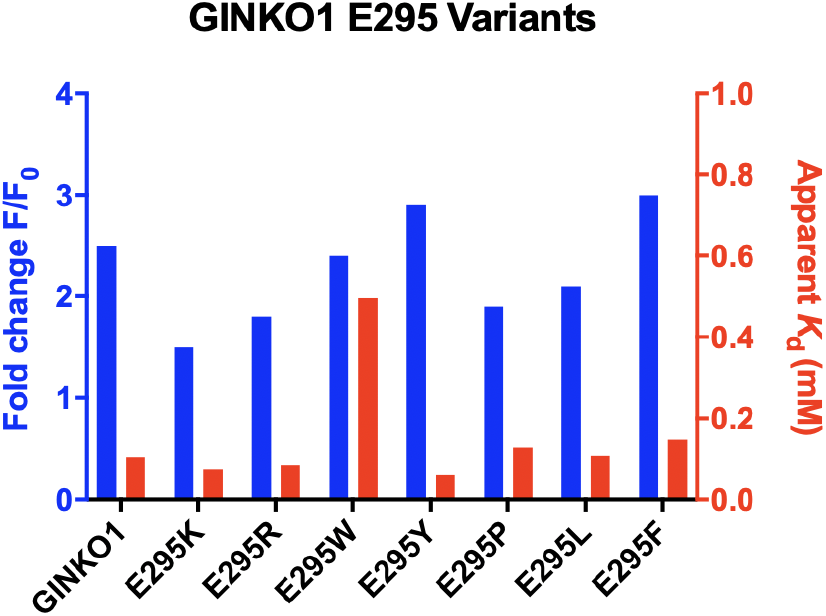
Comparison of E295 variants of GINKO1. The bulky hydrophobic residues Y and F lead to improved F_max_/F_0_. E295W retained the F_max_/F_0_, but simultaneously increased the apparent *K*_d_ substantially. E295K and E295R resulted in a smaller F_max_/F_0_ than template GINKO1.

**Supplementary Figure 4.**
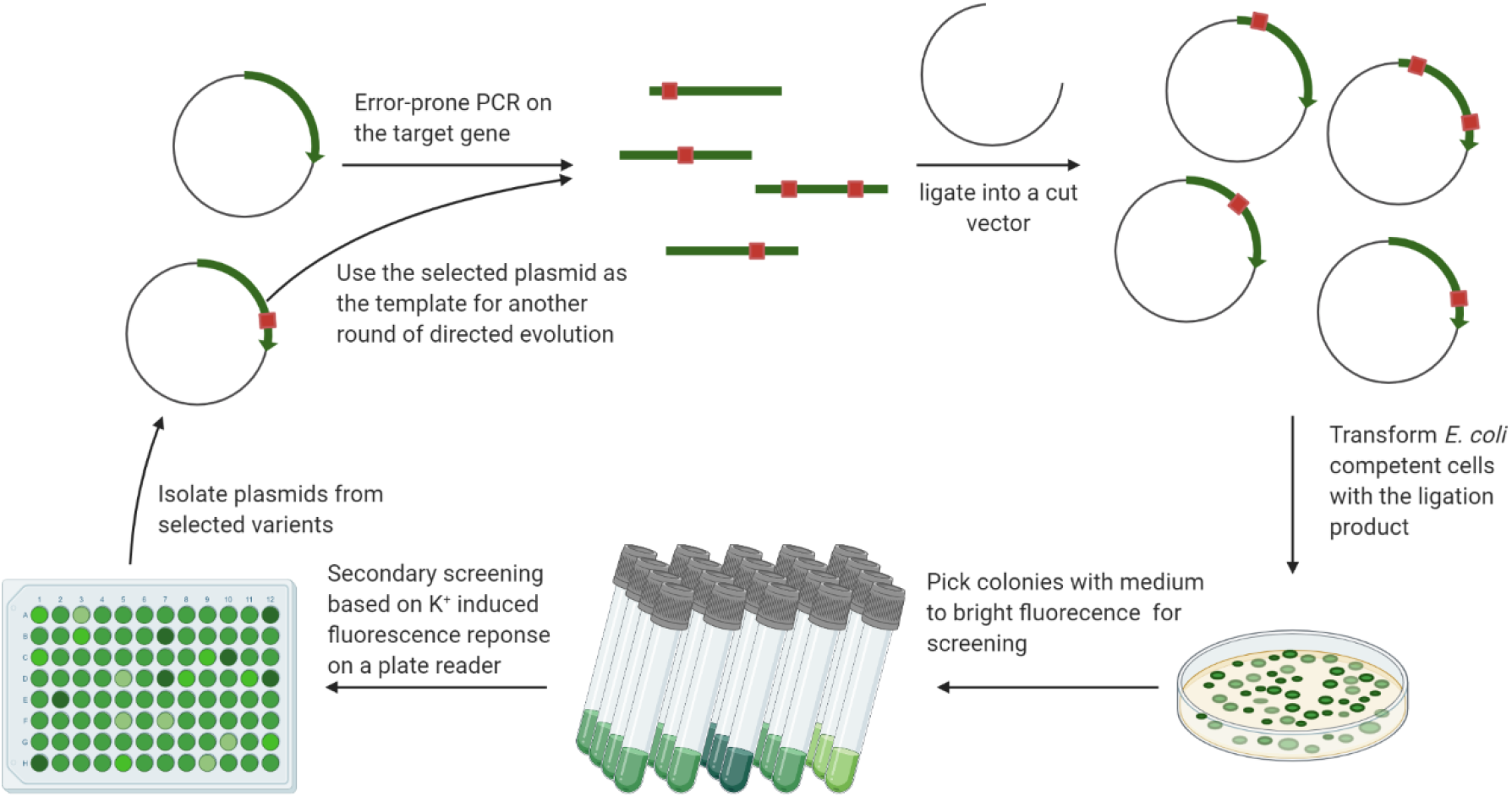
The scheme of directed evolution. Error-prone PCR was used to amplify the GINKO gene with random mutations. The PCR products were digested and ligated into a pBAD vector. After transformation, 5000–10000 colonies were visually inspected and around 40–80 were picked and cultured based on their brightness. The cultures were then pelleted and GINKO variants were extracted via free-and-thaw. The lysates were screened in a plate reader with fluorescence measurements in the absence and presence of K^+^. Variants with the best performance were used as templates for the next iterative round of evolution.

**Supplementary Figure 5.**
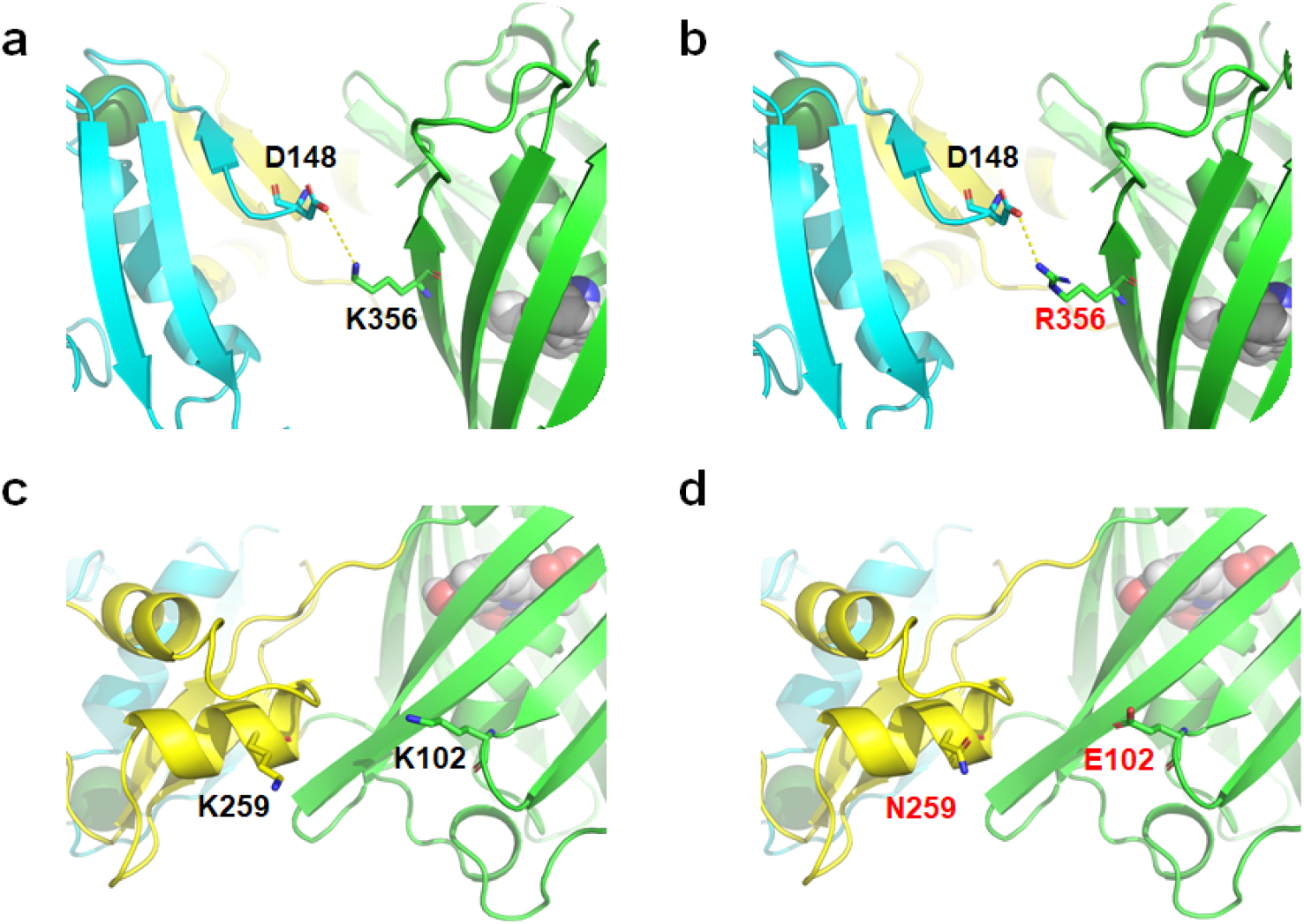
Selected mutations positioned in the GINKO1 structure. K356R (**a** and **b**) led to an improved fluorescence change (F_max_/F_0_) with a possible stronger electrostatic interaction. **a** K356 (green sticks) is in proximity to D148 (cyan sticks) to form electrostatic interaction. **b** R356 (green sticks) side chain is more extended than the K356 side chain, potentially providing a stronger charge attraction with a shorter distance to the D148 side chain. K102E and K259N (**c** and **d**) reduce charge repulsion between two lysine residues. **c** K102 (yellow sticks) and K259 (green sticks) repulse each other with the same charge. **d** E102 (yellow sticks) and N259 (green sticks) removed the charge repulsion, potentially leading to a stable interface between Kbp and EGFP.

**Supplementary Figure 6.**
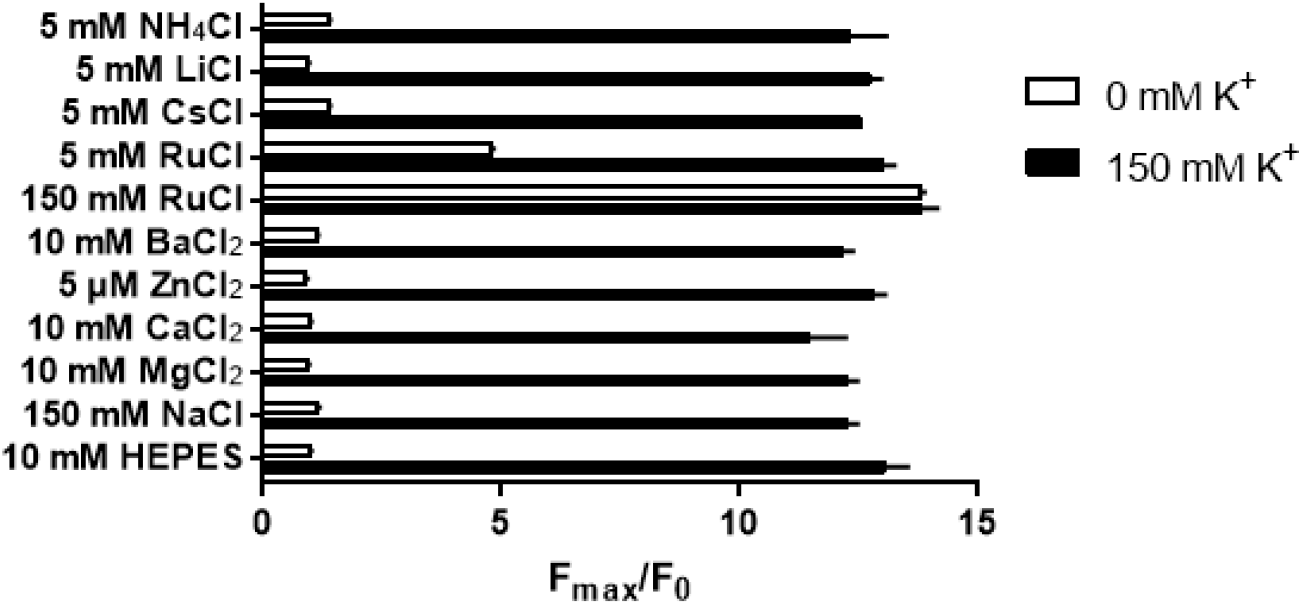
Ion competition assay of GINKO2. To examine whether other cations in the environment could affect the K^+^ response of GINKO2, a specificity test was performed in both 0 and 150 mM K^+^ buffer (n=3). Without K^+^, only Rb^+^ was able to induce a fluorescence change. In presence of 150 mM K^+^, the fluorescence change induced by K^+^ was not affected by any other cations.

**Supplementary Figure 7.**
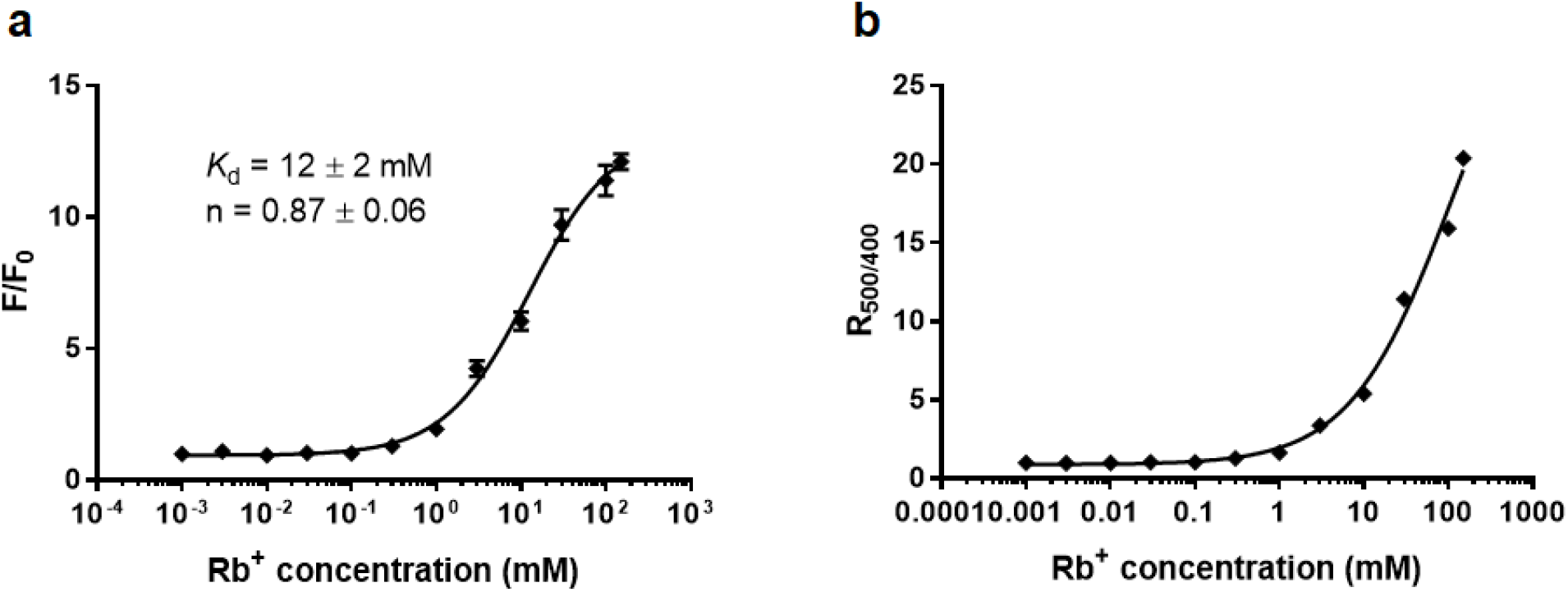
Rb^+^ titration of GINKO2. **a** The fluorescence change (F/F_0_) of GINKO2 *versus* Rb^+^ concentration. n=3. **b** R_500/400_ *versus* Rb^+^ concentration. GINKO2 responds to Rb^+^ ratiometrically.

**Supplementary Figure 8.**
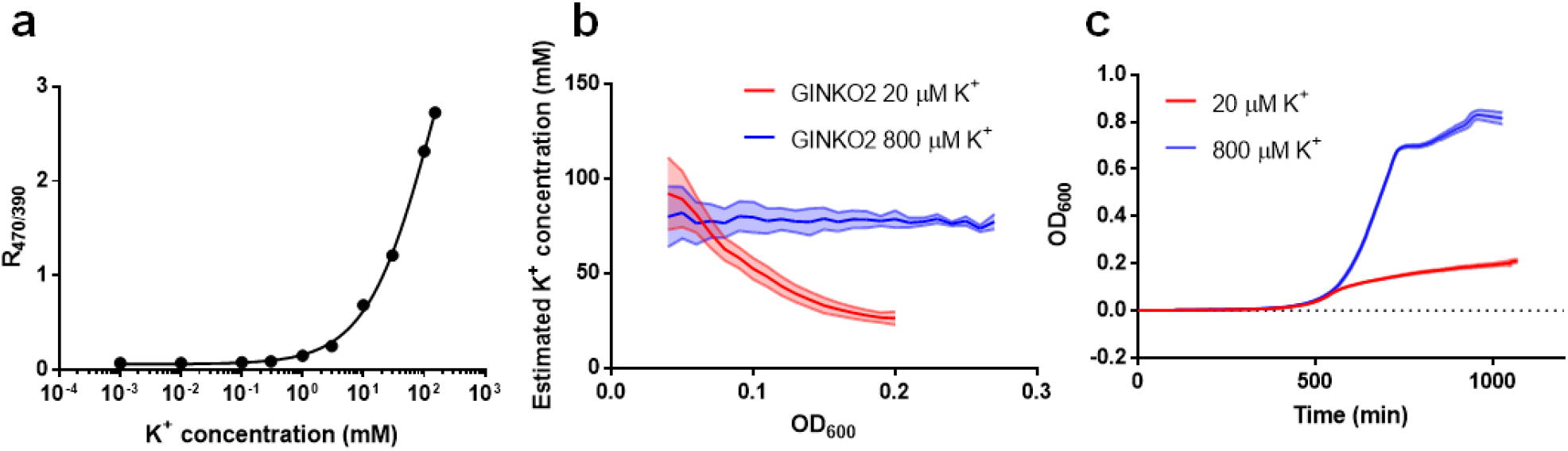
Effect of growth medium K^+^ concentration on *E. coli* intracellular K^+^ concentration and growth rate. **a** Excitation ratio R_470/390_ of purified GINKO2 *versus* K^+^ concentration at pH 7.4. This curve serves as a calibration curve for the estimation of intracellular K^+^ concentrations. **b** Estimated K^+^ concentration change during *E. coli* growth. While *E. coli* grew in an environment with a micromolar level of K^+^, they were able to accumulate K^+^ in a millimolar level. The intracellular K^+^ dropped from 100 mM to 28 mM when *E. coli* grew from OD_600_ of 0.04 to 0.2 in medium supplemented with 20 µM K^+^. The estimated K^+^ concentration remained around 80 mM for cells that grew in 800 µM K^+^. **c** Growth curves of *E. coli* in 20 µM and 800 µM K^+^ medium. These two growth curves indicated that cells growing in the medium supplemented with 20 µM experienced a slower rate of growth, likely due to the limited availability of K^+^.

**Supplementary Figure 9.**
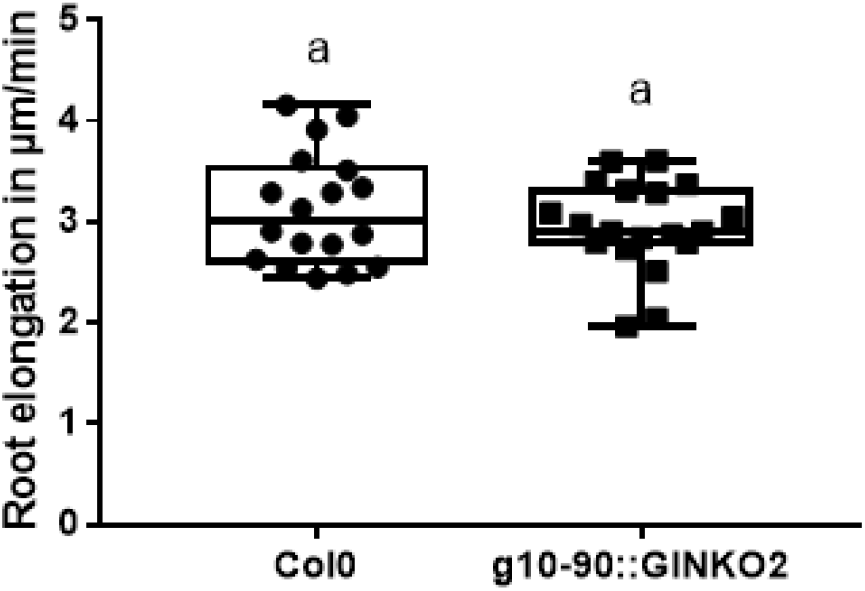
Root elongation of wild type and GINKO2-expressing *Arabidopsis thaliana*. Comparison of root elongation between wild type (WT) Columbia 0 *Arabidopsis thaliana* ecotype to Columbia 0 seedlings expressing g10-90::GINKO2. n = 18 individual seedlings for the Col0 wildtype group, n=19 individual seedlings for the g10-90::GINKO2 group. Letters indicate the significantly different statistical groups with P < 0.05 minimum. Statistical analysis was conducted with a nonparametric multiple comparison

**Supplementary Figure 10.**
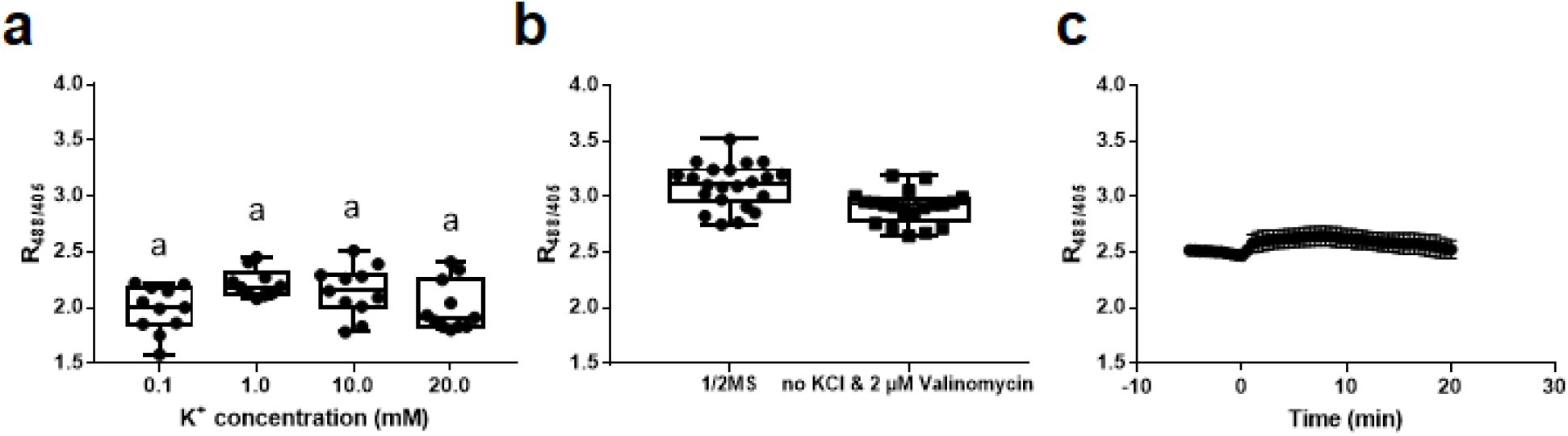
GINKO2 responses in plants under various conditions. **a** Effect of increasing concentrations of KCl on g10-90::GINKO2 R_488/405_ without K^+^ depletion pre-treatment. n ≥ 10 individual seedlings. Letters indicate the significantly different statistical groups with P < 0.05 minimum. Statistical analysis was conducted with nonparametric multiple comparisons. **b** Effect of a six-hour K^+^ depletion treatment with 0 mM KCl and 2 µM valinomycin on g10-90::GINKO2 R_488/405_. 1/2MS: Murashige and Skoog medium half strength. n ≥ 20 individual seedlings. P <0.01. **c** Effect of 100 mM NaCl on g10-90::GINKO2 R_488/405_ in root tip without prior K^+^ depletion. Treatment was applied at time zero. n = 9 individual seedlings.

**Supplementary Figure 11.**
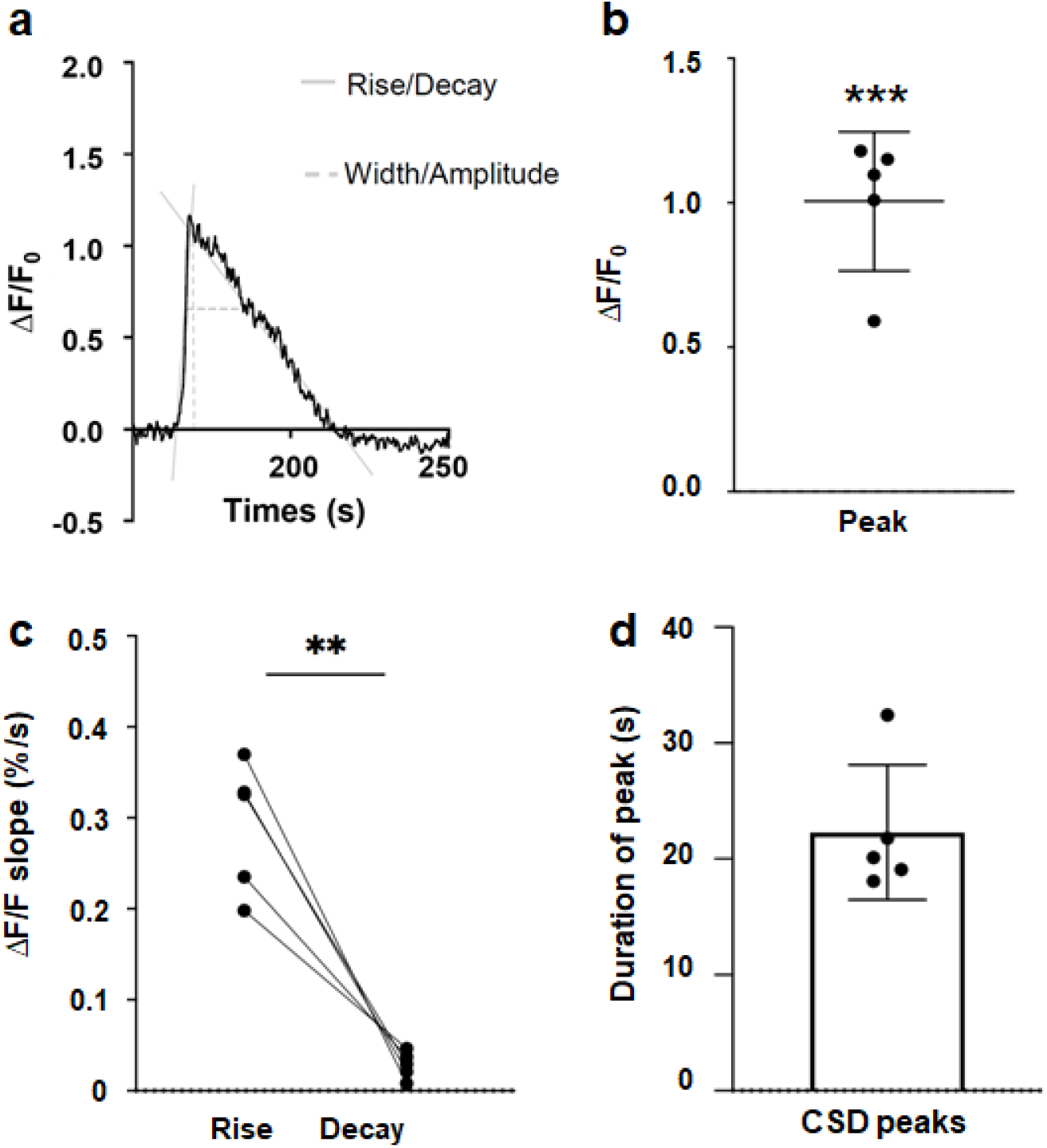
GINKO2 response curve of a representative mouse CSD wave, CSD response amplitude, rise and decay slopes, and CSD duration. **a** An example of a CSD wave showing decay, rise, width, and amplitude. **b** ΔF/F peaks during CSD waves. N = 2 mice, n = 5 ROIs, paired t-test, ***p = 0.0007. **c** Calculated slope coefficient using simple linear regression of the rise and the decay of CSD waves. N = 2 mice, n = 5 ROIs, paired t-test, **p = 0.0024. **d** Average CSD wave duration N = 2, n = 5, mean ± s.d.

**Supplemental Figure 12.**
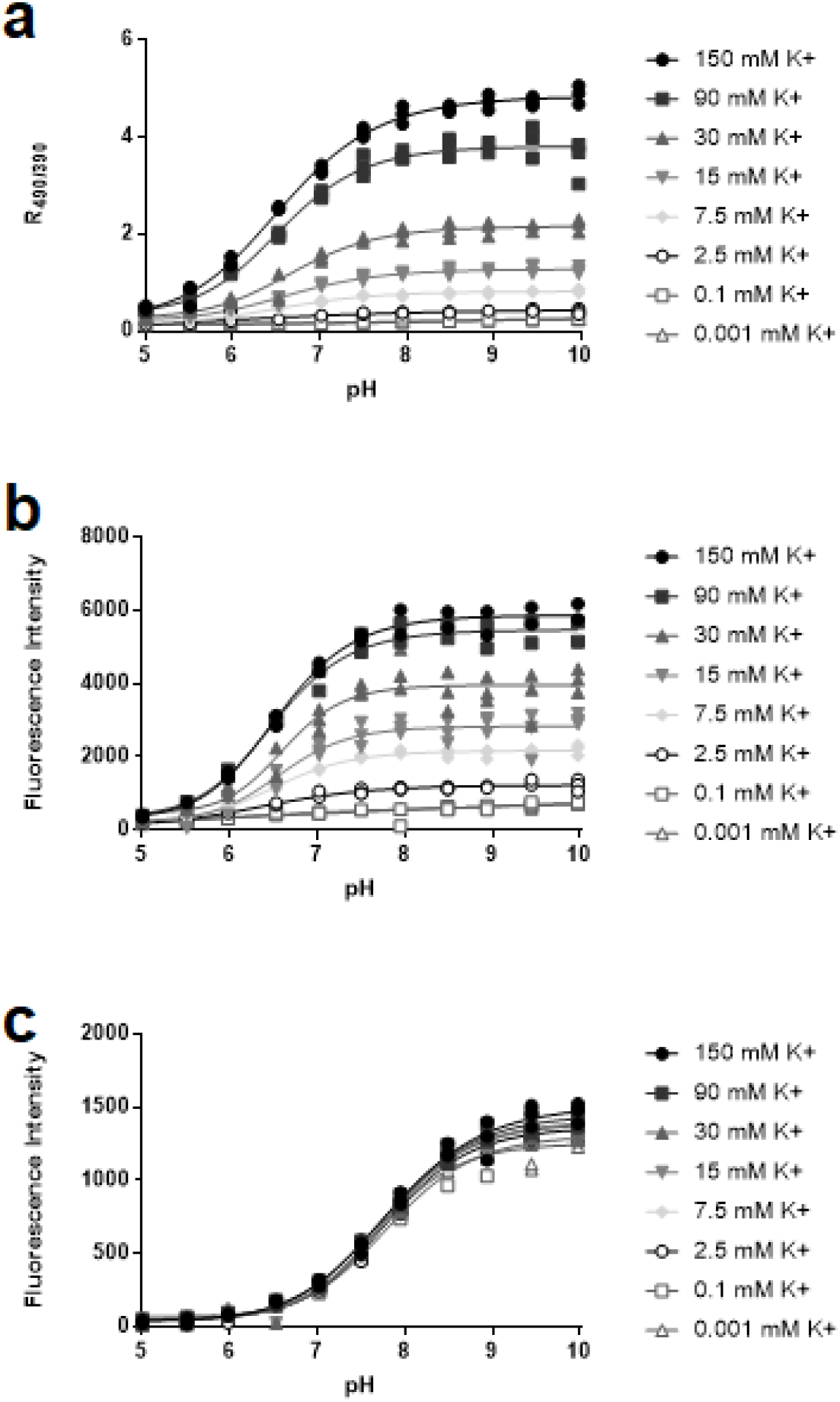
pH titration of pHuji-GINKO2 at various concentrations of K^+^. The titration curves (**a** and **b**) of pHuji-GINKO2 resemble those of GINKO2 (**Fig. 1i** and **Supplementary Fig. 8a**). **a** The excitation ratio R_490/390_ of GINKO2 *versus* pH. **b** The emission maxima of GINKO2 *versus* pH. **c** The emission maxima of pHuji *versus* pH.

**Supplemental Figure 13.**
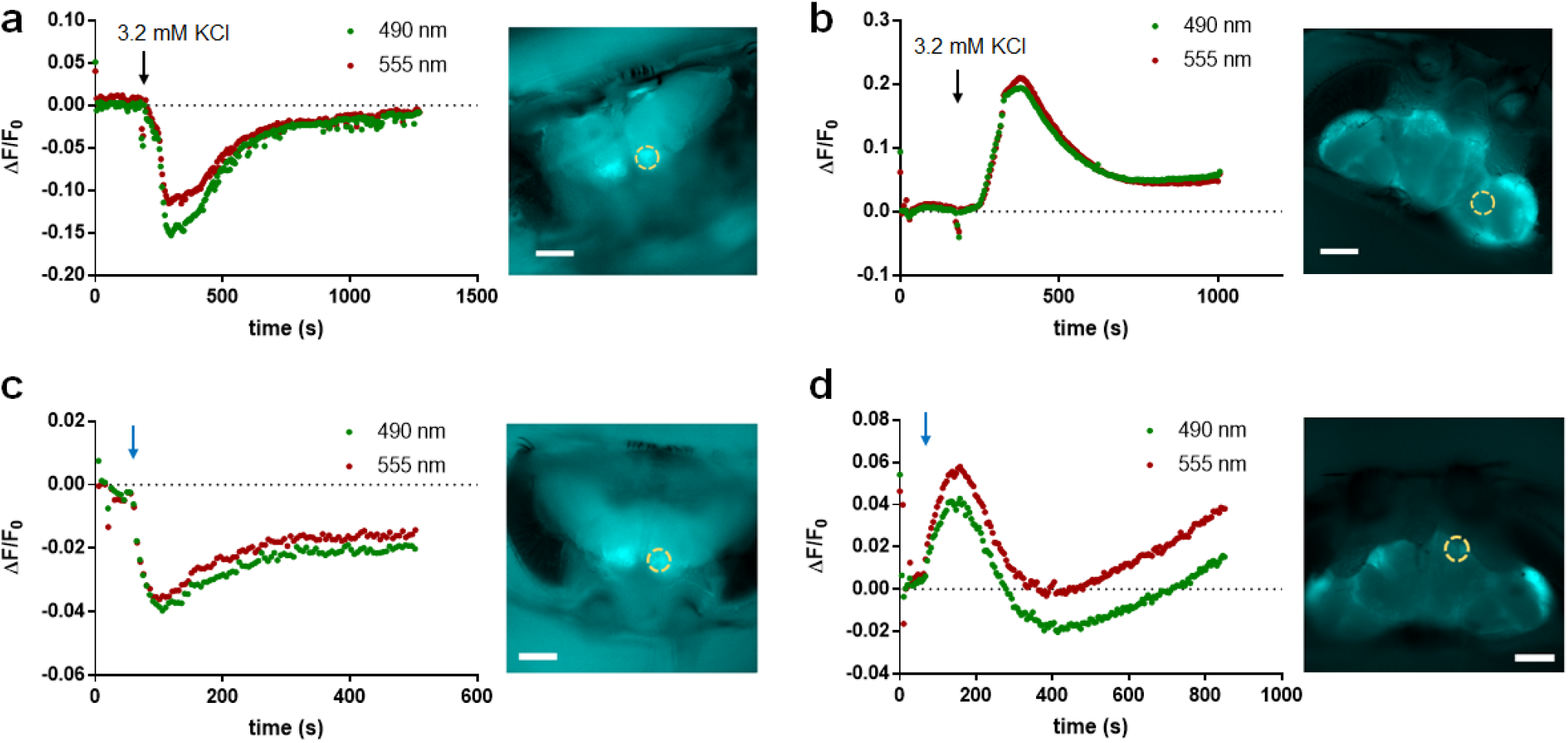
pHuji-GINKO2 responses in the *Drosophila* brain under stimuli. Fly brains expressing pHuji-GINKO2 in (**a**) neurons and (**b**) glia were stimulated by 3.2 mM KCl. The black arrow indicates the time at which 3.2 mM KCl was added. Fly brains expressing pHuji-GINKO2 in (**c**) neurons and (**d**) glia were stimulated by 500 impulses electrical stimuli at 50 Hz, at the time indicated by the blue arrow, by a glass electrode. Scale bars: 100 µm.

## Supplementary Note

### GINKO1 crystallization and structure comparison with Kbp

To aid our protein engineering effort, we obtained crystals of GINKO1 in the buffer supplemented with 150 mM K^+^ (**Supplementary Fig. 1, Supplementary Table 1**). Previous work by Ashraf and colleagues suggested challenges in crystallization of Kbp for X-ray crystallography, which led to an alternative approach to obtain an NMR solution structure of Kbp^10^. We suspect that fusing Kbp to EGFP constrains the conformational mobility of Kbp, thus increasing the stability of Kbp protein for it to be crystallized as a domain in GINKO1. A similar approach has been reported recently to stabilize small transmembrane proteins for crystallization^36^. The Kbp region of the GINKO1 structure aligns well with the previous Kbp NMR solution structure (**Supplementary Fig. 2**). The BON domain and the LysM domain of Kbp were both well-resolved in the GINKO1 structure. The structure further revealed that the K^+^ binding site is located in the BON domain, close to the interface between the BON and LysM domains (**Supplementary Fig. 2**), consistent with the previous work by Ashraf and coworkers who suggested that the BON domain binds K^+^ and the LysM domain stabilizes the K^+^-bound BON domain^10^.

### Structural analysis of selected mutations

A total of 14 amino acid mutations were accumulated in GINKO2 engineering (**Supplementary Table 2**). With the structural insights, we were able to rationalize some critical mutations. The K356R mutation doubled the F_max_/F_0_. In the GINKO1 structure, K356 is located at the interface of the Kbp and GFP (**Supplementary Fig. 5a**). K356R likely allows the K^+^-bound GINKO to be more stable via a shorter distance to and a stronger charge interaction with D148 (**Supplementary Fig. 5b**), which may subsequently improve the F_max_/F_0_. Another case is mutations on a pair of lysines facing each other: The K259N and K102E first appeared in two different variants in the GINKO1.5 library and provided small improvements. GINKO1 structure revealed K259 and K102 reside on Kbp and GFP respectively (**Supplementary Fig. 5c**). When using both variants as templates to generate the GINKO1.6 library, K259N and K102E were simultaneously incorporated in the selected GINKO1.6.15 variant, which provided a substantial improvement of F_max_/F_0_ from ∼3.5 to 4.5. This mutation combination likely removes the repulsive force and stabilizes the interaction between Kbp and GFP (**Supplementary Fig. 5d**).

### K^+^ depletion reduces the buffering effect from the vacuolar K^+^ reservoir

Vacuoles are K^+^ reservoirs with up to 200 mM of K^+^. The significant stock of vacuolar K^+^ is available to the cytoplasm for the overall regulation of cytoplasmic K^+^ concentration^37^. Due to the extremely low vacuolar pH, GINKO2 fluorescence is not visible in vacuoles thus can not be used to report vacuolar K^+^ concentration change. When we transferred the seedlings from the MS medium with 19 mM K^+^ to K^+^ gradient buffers (0.1, 1, 10, and 20 mM) for 2.5 days, cytosolic GINKO2 fluorescence reported no significant differences in R_488/405_ across the concentration range (**Supplementary Fig. 10a**). We reasoned that the vacuolar pool of K^+^ regulates cytosolic K^+^ homeostasis, buffering the low K^+^ in the treatments. It was previously reported that during low K^+^ treatment, the vacuolar pool of K^+^ gradually decreases to sustain the cytosolic pool, and only when the vacuolar pool is severely diminished that the cytosolic K^+^ concentration starts to decline^37^. Therefore, we thought to deplete the vacuolar K^+^ before imaging to reduce its buffering effect, by transferring the seedlings onto a medium containing 0 mM K^+^ and the K^+^-specific ionophore valinomycin (2 µM). This pre-depletion of K^+^ enabled direct manipulation of the cytosolic K^+^ concentration using media of different K^+^ concentrations, allowing GINKO2 to display its full sensing capacity (**Fig. 2c**). We observed a significant decrease of the GINKO2 R_488/405_, indicating a lowered concentration of cytoplasmic K^+^ (**Supplementary Fig. 10b**). The NaCl treatment without pre-depletion of K^+^ produced an initial increase in cytoplasmic K^+^ concentration followed by a decrease after 10 minutes (**Supplementary Fig. 10c**). This, again, can be attributed to the vacuolar K^+^ exporting into the cytoplasm. With K^+^ pre-depletion, we observed a more pronounced decrease of GINKO2 R_488/405_ in plants (**Fig. 2d, 2e**). Taken together, our results with cytoplasmic GINKO2 reflected the complexity of the regulatory mechanism in plants, therefore caution should be applied when investigating K^+^ dynamics in tissues or zones with different vacuole sizes (e.g., epidermis *versus* stele, dividing *versus* elongating cells). Nonetheless, GINKO2 demonstrated great sensitivity in reporting K^+^ changes in the roots of *Arabidopsis thaliana*. With appropriate protocols, GINKO2 represents a great step forward for the study of K^+^ homeostasis in plants and could be used to detect phenotypes in mutants (e.g. in K^+^ transporters) and characterize detailed K^+^ dynamics under disturbing stresses.

### Changes is pH may distort estimates of K^+^ changes

Fluorescent protein-based biosensors are often pH-sensitive^38,39^, which brings complications to applications where the pH changes, restricting their applicability. The p*K*_a_ of GINKO2 (6.8) is very close to physiological pH, thus changes in pH during an experiment could induce fluorescence intensity changes that can be confused with those induced by K^+^.

The pH sensitivity of GINKO2 was taken into careful consideration in the applications in bacteria, plants, and mice demonstrated in the main text (**Fig. 2**). In the *E. coli* growth experiment, a pH control pHluorin was used to monitor pH changes and indicated a well-maintained intracellular pH during the experiment (**Fig. 2a**). In plant cells, previous research established that cytosolic pH is tightly regulated and well-maintained^40^. More importantly, intracellular pH remained steady in *Arabidopsis thaliana* root cells after the NaCl treatment, as demonstrated by previous study^41^ and in our own control experiment with pHGFP (**Fig. 2e**, lower panel). The fluorescence increases observed with GINKO2 during CSD (**Fig. 2h**) corresponds well to descriptions of extracellular K^+^ concentration dynamics previously reported during CSD^22^. Furthermore, the dynamic changes in pH during CSD (short increase in pH ∼5 s, followed decrease in pH^42^) does not correspond to the fluorescence dynamics we observed with GINKO2, strongly indicating that the fluorescence dynamics of GINKO2 observed during CSD corresponds to changes in extracellular K^+^ concentration. Overall, GINKO2 is well poised for these applications as pH remained constant or resulted in a GINKO2 signal change in the opposite direction to that caused by K^+^. Yet, caution must be exercised when applying GINKO2 under unknown pH dynamics.

In our attempts to visualize potential K^+^ changes *in vivo* in *Drosophila*, we combined GINKO2 with a red pH biosensor, pHuji^43^, to monitor both K^+^ and pH concurrently. We first characterized pHuji-GINKO2 fusion protein *in vitro*. As shown in **Supplementary Fig. 12**, decreasing pH reduces the fluorescence of GINKO2, but does not change the affinity for K^+^. In addition, the fluorescence of pHuji is not sensitive to the K^+^ concentration. We then produced transgenic flies expressing pHuji-GINKO2 under control of the Gal4-UAS system, either in neurons (elav-Gal4) or glia (repo-Gal4). Fly brains were stimulated either by rapidly elevating the extracellular K^+^ concentration, or electrically with a glass electrode. In neurons, both stimuli led to a decline in GINKO2 fluorescence while in glia electrical stimulation and the application of high K^+^ led to an increase in GINKO2 fluorescence. However, these stimuli also led to similar changes in pHuji fluorescence, indicating substantial pH changes (**Supplementary Fig. 13**). Although it is expected that stimulated neuronal activities are likely to lead to the efflux of K^+^ and has been observed by others in several different preparations^44^, due to the susceptibility of GINKO2 to pH interference, the GINKO2 fluorescence change observed in this particular set of experiments can not be conclusively interpreted as K^+^ changes in the simulated neurons or glial cells.

The pH sensitivity of GINKO2 presents challenges in applications with potentially unknown pH changes. pHuji-GINKO2 offers a way to monitor such pH changes thus avoiding misinterpretation of GINKO2 fluorescence change. To further address this challenge, future efforts in biosensor development should be directed to engineer GINKO with less pH-sensitive FP or optimize GINKO to achieve a lower p*K*_a_. This should lead to GINKO variants with reduced pH sensitivity for a broader range of applications at the physiological pH.

## Supplementary Methods

### *In vivo* imaging in *Drosophila*

To generate transgenic flies expressing pHuji-GINKO2 under the control of the Gal4-UAS system, pHuji-GINKO2 was cloned into the pUAST vector^45^. The vector was injected into *w*^*1118*^ embryos (BestGene, Inc.) and transformant lines with insertions on each major chromosome were selected. To drive expression in all neurons, UAS-pHuji-GINKO2 flies were crossed to *w*^*1118*^ *elav-Gal4*^*C155*^. To drive expression in glia, UAS-pHuji-GINKO2 flies were crossed to *w*^*1118*^ *repo-Gal4/TM3, Sb*. The head capsules of flies were opened using the goggatomy procedure^46^, where the head is rapidly encapsulated in a photopolymerizable resin and then sliced to expose the live brain. Heads were cut transversely along a line through the joints between the second and third antennal segments. All experiments were performed in saline with the following composition: 120 mM NaCl, 3 mM KCl, 1 mM CaCl_2_, 4 mM MgCl_2_, 4 mM NaHCO_3_, 1 mM NaH_2_PO_4_, 8 mM D-trehalose, 5 mM D-glucose and 5 mM TES (pH 7.2). The bath solution (∼2.5 mL) was oxygenated and stirred by directing an airstream over the solution. Glass electrodes filled with saline were used to stimulate the brain and timing was controlled by an A.M.P.I. Master-8 (Microprobes for Life Science). Fly brains were imaged on a BX50WI upright microscope (Olympus) with an ORCA-Flash 4.0 CMOS camera (Hamamatsu). GINKO2 fluorescence was monitored at 510 nm with excitation at 402 and 490 nm, whereas pHuji was excited at 555 nm and the emission was monitored at 610 nm. Illumination was provided by a LED (CoolLED) through a Pinkel filter set (89400 - ET - DAPI/FITC/TRITC/Cy5 Quad, Chroma). Images were acquired with MetaMorph software (Molecular Devices).

